# Cognitive dysfunction following brain trauma results from sex-specific reactivation of the developmental pruning processes

**DOI:** 10.1101/2024.08.13.607610

**Authors:** Dena Arizanovska, Carlos A. Dallera, Oluwarotimi O. Folorunso, Gerald F. Bush, Jennifer B. Frye, Kristian P. Doyle, Jonathan R. Jagid, Herman Wolosker, Bernardo A. Monaco, Joacir Graciolli Cordeiro, Coleen M. Atkins, Anthony J. Griswold, Daniel J. Liebl

## Abstract

Cognitive losses resulting from severe brain trauma have long been associated with the focal region of tissue damage, leading to devastating functional impairment. For decades, researchers have focused on the sequelae of cellular alterations that exist within the perilesional tissues; however, few clinical trials have been successful. Here, we employed a mouse brain injury model that resulted in expansive synaptic damage to regions outside the focal injury. Our findings demonstrate that synaptic damage results from the prolonged increase in D-serine release from activated microglia and astrocytes, which leads to hyperactivation of perisynaptic NMDARs, tagging of damaged synapses by complement components, and the reactivation of developmental pruning processes. We show that this mechanistic pathway is reversible at several stages within a prolonged and progressive period of synaptic loss. Importantly, these key factors are present in acutely injured brain tissue acquired from patients with brain injury, which supports a therapeutic neuroprotective strategy.

## INTRODUCTION

Despite affecting nearly 70 million people each year, few treatments are currently available for traumatic brain injured (TBI) patients^1^. This is largely due to a poor mechanistic understanding of the complex underlying pathologies, in which both the primary and secondary injury phases play a pivotal role^2^. Furthermore, the unique nature of each injury, which varies from mild to severe, makes it difficult to apply the same therapeutic principles across patients. One pathological aspect of TBI preserved across injury types is diffuse synaptic damage, leading to broad alterations in synaptic transmission and plasticity, and often underlying long-term cognitive impairment^3–7^.

We have previously shown that glial D-serine contributes to synaptic damage in the hippocampus, a brain region critical for learning and memory, using a controlled cortical impact (CCI) injury mouse model^6,7^. D-serine, the predominant co-agonist of N-methyl-D-aspartate receptors (NMDARs) in cortico-limbic areas, is converted from L-serine by the serine racemase (SRR) enzyme^8–11^. Under physiological conditions, SRR is predominantly expressed in neurons; however, its expression is increased in both astrocytes and microglia after CCI injury^6,7,12,13^. Blocking D-serine synthesis or release from either glial cell type is sufficient to prevent hippocampal synaptic damage and cognitive dysfunction after CCI injury in male mice^6,7^. The mechanism of glial D-serine-mediated damage and its role in female mice have not been evaluated.

Many studies have reported different outcomes based on biological sex in animal models and human patients, where estrogen is thought to play a neuroprotective role^14–21^. Moreover, there are sex differences in microglial activation after injury and under other neuroinflammatory conditions^16,22–25^, raising the possibility that microglial D-serine responses might be differentially regulated between sexes after TBI. Here, we investigated the mechanism of glial D-serine-induced synaptic damage after TBI and in both male and female mice. We examine whether tonic glial D-serine release leads to sublethal hyperactivation of GluN2B-containing perisynaptic NMDARs, causing synaptic damage in the absence of cell death. Hyperactivation of GluN2B-NMDARs has been associated with decreased synaptic plasticity^[26]^, dendritic spine damage in models of cerebral ischemia^26^, Alzheimer’s Disease^27^, and Huntington’s Disease^28^. Our studies also link pathological D-serine activation of GluN2B-NMDARs to the reactivation of developmental synaptic pruning processes, including phosphatidylserine (PS) externalization and complement-mediated tagging of synapses. In addition, we compared these processes in male and female mice and examined whether perilesional tissues from TBI patients undergo similar processes. Together, these studies aim to elucidate sex-dependent outcomes as well as mechanisms of synaptic damage after injury with the goal of contributing to more targeted treatments for TBI patients.

## RESULTS

### Female mice have less glial serine racemase expression, synaptic damage, and cognitive dysfunction than male mice after CCI injury

Under physiological conditions, serine racemase (SRR) is predominantly expressed by post-synaptic neurons and converts L-serine to D-serine, which in turn activates NMDARs to facilitate synaptic plasticity^12^ (Figure 1A). Following TBI, astrocytes and microglia upregulate SRR expression, leading to increased and prolonged D-serine synthesis and release. D-serine from both glial cell types is necessary to reach threshold levels to induce synaptic damage and cognitive dysfunction in male mice^6,7^ (A). To examine whether there were sex differences in our model, we first performed contextual fear conditioning on male and female WT mice. We observed a significant one-third (34%) reduction in freezing behavior compared with sham controls in male but not female mice at 7 days post-injury (dpi) (Figure 1B). To determine whether the observed sex differences in cognition were due to alterations in hippocampal astrocytic or microglial expression of the rate-limiting enzyme SRR, we used flow cytometry to determine the cell-specific SRR levels. Microglia were identified as CD11b^+^/CD45^low^ labelled viable cells (Figure S1C-F) and astrocytes were identified as CD11b^-^/CD45^-^/ASCA2^+^ viable cells (Figure S1G-J), as previously validated^7^. Positive SRR signals were determined against secondary-only staining (Figure S1A-B,1K-R). Astrocytes showed a significant 10-20% increase in SRR-positive cells in both males and females (Figure 1C). However, only male mice showed a significant 30% increase in microglia expressing SRR at 7 dpi, which was 2-fold greater than that in female microglia (Figure 1D). Furthermore, male but not female mice had a significant ∼20% increase in mean fluorescent intensity (MFI) per microglia and astrocytes at 1 or 7 dpi (Figure S2A-D). These results are supported by immunohistochemistry (IHC), where SRR was expressed at 7 dpi (Figure 1E-F) and colocalized with GFAP+ astrocytes in the CA1 region of male (Figure 1G) and female (Figure 1I) hippocampi. Similar to the flow cytometry results, IHC revealed colocalization of SRR with IBA1+ microglia to a greater extent in males (Figure 1H) than in females (Figure 1J) at 7 dpi.

**Figure 1.**
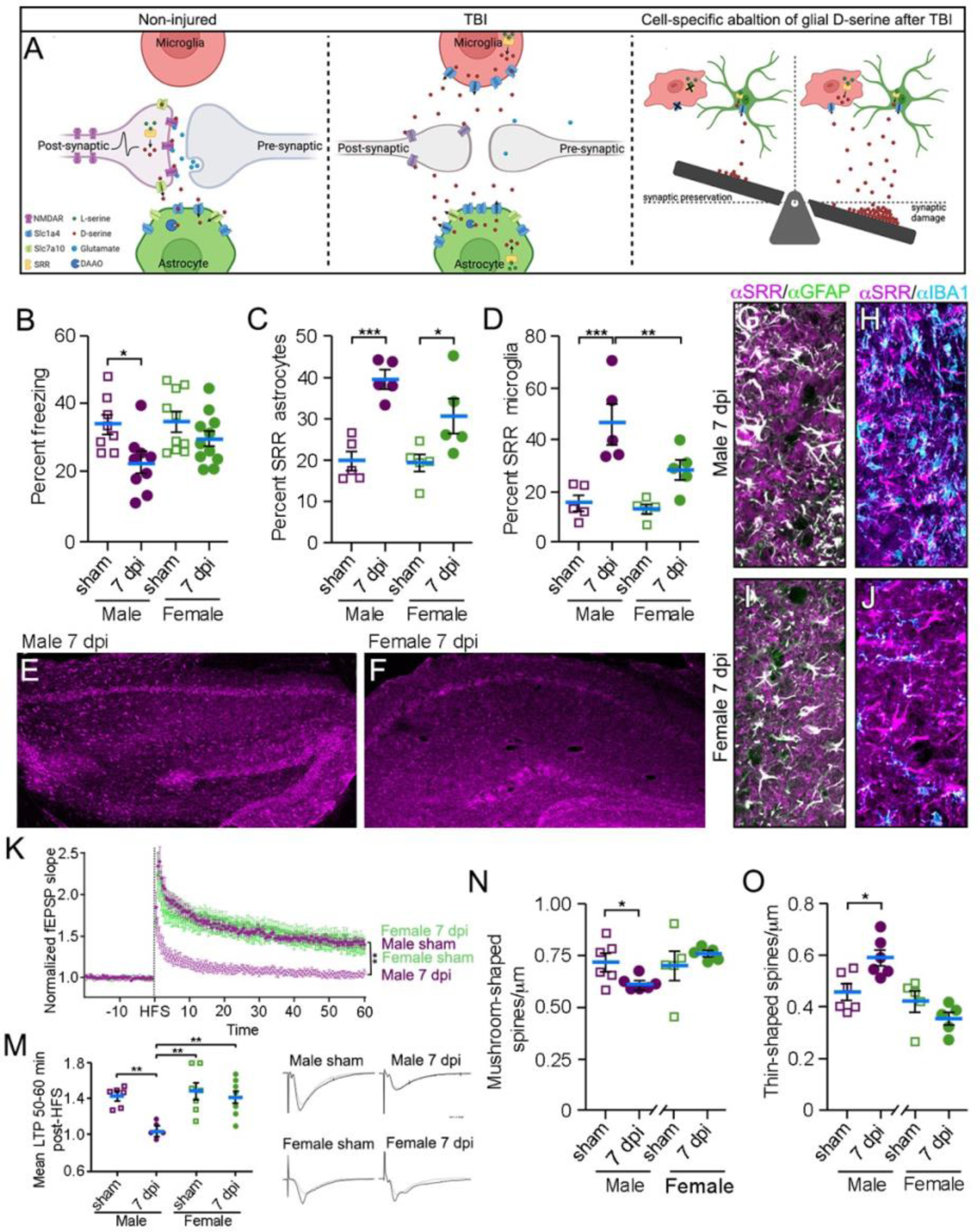
Reduced CCI-induced microglial SRR and synaptic damage in female mice. (A) Schematic of D-serine (red dots) production in homeostatic non-injured and pathological conditions (created with Biorender). Preventing D-serine release from either glial cell type is sufficient to rescue CCI-induced synaptic damage and cognitive deficits. (B) CCI induces freezing deficits in male but not female WT mice at 7 dpi. (C) Flow cytometry indicates increased SRR expression in male and female astrocytes at 7 dpi. (D) Flow cytometry indicates reduced SRR expression in female compared to male microglia at 7 dpi. (E-F) SRR staining patterns in the hippocampus of male and female CCI-injured mice that colocalize with (G, I) GFAP+ astrocytes and (H,J) IBA1+ microglia. (K,M) Electrophysiological LTP recordings in male and female mice at 7 dpi. (N) Golgi staining shows losses in mature mushroom shaped spines and (O) increases in degenerating thin spines in male but not female mice at 7 dpi. (K) Scale bar: 1 μM. *p<0.05, **p<0.01, ***p<0.001. (B-D) Two-way ANOVA with Tukey’s multiple comparison test; (K,M) Repeated measures two-way ANOVA. (B) n=8-11, (C,D) n=5, (M) n=5-7. (N,O) Student’s t-test, n=5-6. Values represent mean ± SEM.

Given the differences in cognitive dysfunction and SRR expression, we next examined field potentials in the CA1 *stratum radiatum* in response to high-frequency electrical stimulation (HFS) of the Schaffer collaterals in male and female mouse hippocampal slices (Figure 1K,M). Following the delivery of a monophasic electrical impulse, input-output (I/O) curves indicated reduced field excitatory post-synaptic potentials (fEPSPs) at 7 dpi in male mice, but not in females, compared with their respective sham controls (Figure S2G). No differences were observed between male and female mice using paired-pulse facilitation (PPF) (Figure S2H). Potentiation of synaptic transmission following tetanic stimulation to induce long-term potentiation (LTP) was only impaired in male mice at 7 dpi and not in female mice (Figure 1K,M), supporting our conclusion that CCI injury-induced damage was spared in female mice.

Golgi-Cox staining^7,29^ (Figure S2E) was used to evaluate sex differences in the density and morphology of CA1 apical dendrites after CCI injury. There were no changes in the overall spine density at 7 dpi (Figure S2F); however, male but not female mice showed a reduction in mushroom-shaped excitatory spines (Figure 1N) and an upregulation of thin-shaped degenerating spines (Figure 1O). This may underlie the functional learning and memory deficits in male mice (Figure 1B), as mushroom-shaped spines are representative of mature spines that can form functional synapses, whereas thin-shaped spines likely indicate spine degeneration after CCI^7,29^.

To demonstrate that microglial D-serine plays a direct role in synaptic damage after TBI in mice, we employed genetically modified *Tmem119^creERT2^:SRR^fl/fl^* and *SRR^fl/fl^* control mice, where SRR was specifically knocked down in microglial cells (Figure S3). Similar to WT mice, we observed no changes in hippocampal dendritic spine density or morphology (Figure S4A-C) in *SRR^fl/fl^* or *Tmem119^creERT2^:SRR^fl/fl^* female mice at 7 dpi. This corresponded to no functional deficits in contextual or cued fear-conditioning behavior (Figure S4D,E). Furthermore, sex differences were not due to differences in baseline anxiety (Figure S4F) or locomotion (Figure S4G-I). To explore the ways in which microglial D-serine leads to hippocampal dysfunction in male mice, we performed multiplex assays to evaluate the levels of inflammatory molecules and matrix metalloproteinases (MMPs) at 3 dpi (Figure S5). Interestingly, we observed little change in inflammation in the hippocampus at 3 dpi in our CCI model (Figure S5A-M), except for cytokine IL-13, which was significantly decreased by 45% in control *SRR^fl/fl^*but not in *Tmem119^creERT2^:SRR^fl/fl^* mice at 3 dpi (Figure S5F). This suggests that synaptic damage is independent of inflammation. Conversely, the expression of all MMPs tested were 1.5-2.8-fold higher at 3 dpi in *Tmem119^creERT2^:SRR^fl/fl^* mice than in *SRR^fl/fl^* mice (Figure S5N-R). As both MMPs and IL-13 have been implicated in synaptic plasticity^30,31^, this study provides an avenue for future studies investigating microglial D-serine after injury.

### D-serine modulates NMDAR receptor subunits that lead to changes in dendritic spine density and morphology as well as contextual learning deficits

NMDARs are composed of at least two subunits, GluN1 and GluN2, where GluN2A and GluN2B are predominant in the hippocampus^32^. Although there is no clear delineation, NMDARs at synaptic sites are enriched in GluN2A, whereas perisynaptic NMDARs are enriched in GluN2B (Figure 2A) and are involved in deleterious signaling when hyperactivated^33,34^. To determine whether microglial D-serine modulates NMDAR subunit expression, we performed Western blot analysis of whole hippocampal tissue from sham, 1, and 3 dpi male *SRR^fl/fl^* and *Tmem119^creERT2^:SRR^fl/fl^*mice, where 1 dpi represents a period when significant synaptic damage occurs after CCI. We observed significant increases in NR1 (Figure S6A; Figure S7), GluN2A (Figure 2B,D; Figure S7), and GluN2B (Figure 2C, D; Figure S7) expression in control mice at 1 dpi, which was not present in *Tmem119^creERT2^:SRR^fl/fl^*mice. Next, we examined the phosphorylation levels of GluN2B (pGluN2B) at the Ser^1303^ site, as this can enhance NMDAR conductance and stability in the plasma membrane, contributing to spinal damage^26–28^. While the ratio of pGluN2B to total GluN2B was unchanged after injury (Figure S6B, Figure S7), there was significantly greater pGluN2B expression when normalized to total protein at 1 dpi in *SRR^fl/fl^* mice than in *Tmem119^creERT2^:SRR^fl/fl^* mice (Figure 2E, G; Figure S7). This increase is delayed until 3 dpi in *Tmem119^creERT2^:SRR^fl/fl^* mice, a time point at which synaptic losses have already been observed^7^. Death associated protein kinase 1 (DAPK1), the enzyme that phosphorylates GluN2B, mirrors the pGluN2B expression pattern (Figure 2F, G; Figure S7). Furthermore, the phosphorylation of cAMP response binding element protein (CREB), which is activated by GluN2A and inhibited by GluN2B signaling^34^ (Figure 2J), is increased only in *Tmem119^creERT2^:SRR^fl/fl^*mice after injury (Figure 2H, J; Figure S6C, D; Figure S7). Increased pCREB can led to the production of pro-synaptic factors such as brain derived neurotrophic factor (BDNF), which is altered at 1 dpi in mutant as opposed to 3 dpi in control *SRR^fl/fl^* mice (Figure 2I, J; Figure S7). In addition, pERK1/2, which plays a dual role in pro-survival and pro-death signaling as well as in glial inflammation^35^, is significantly increased at 1 dpi only in control *SRR^fl/fl^* mice (Figure S6E).

**Figure 2.**
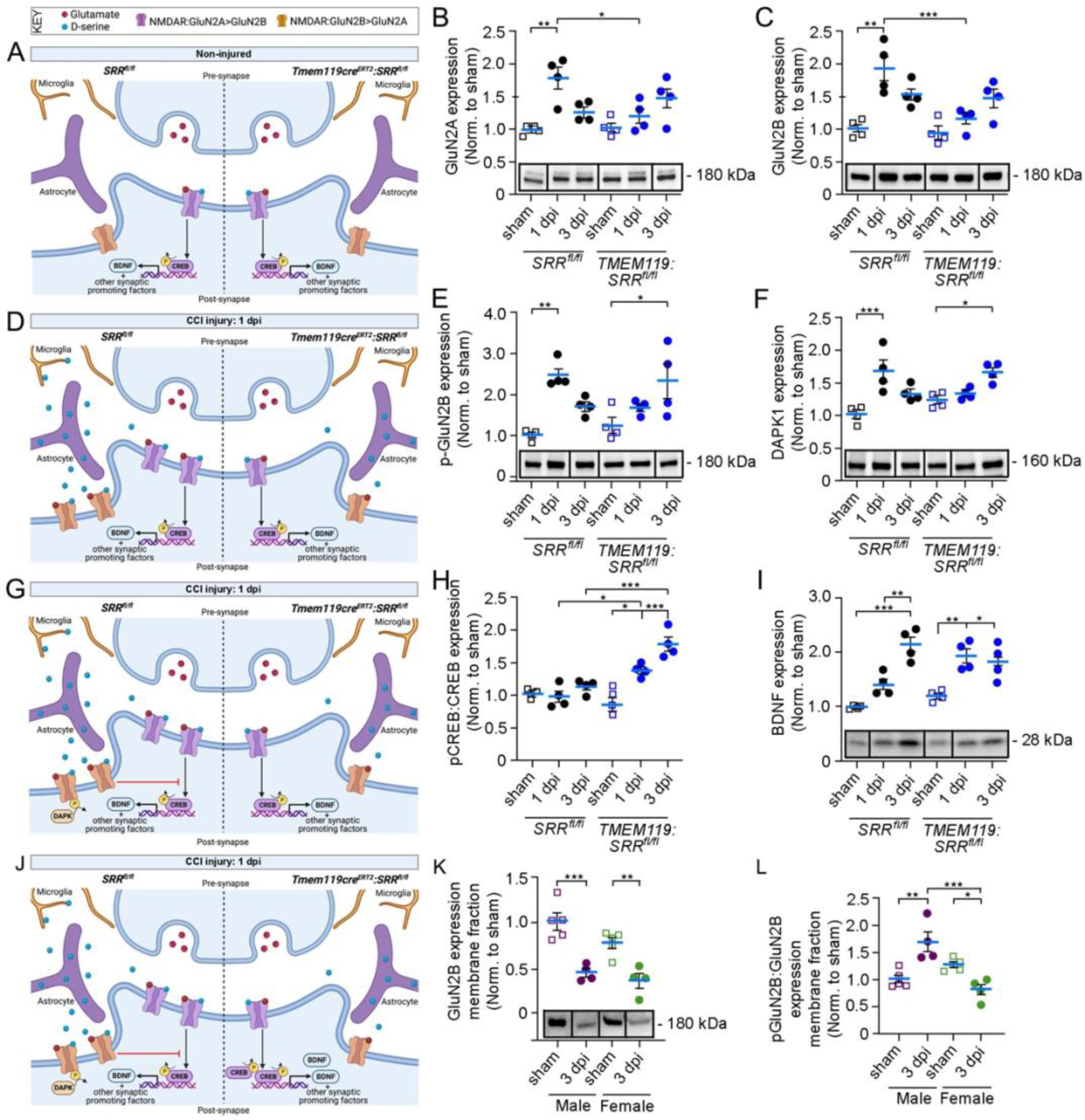
Microglial D-serine alters expression of NMDAR signaling components after injury in male mice. (A, D, G, J) Schematic illustrations in the non-injured quadpartite hippocampal synapse and at 1 dpi of *SRR^fl/fl^* and *TMEM119^creERT2^:SRR^fl/fl^*mice (created with Biorender). (B-D) Western blot of whole hippocampus shows a significant increase in (B) GluN2A and (C) GluN2B expression at 1 dpi in *SRR^fl/fl^* male mice that is not present in microglial knockout mice. (E-G) Expression of (E) phosphorylated GluN2B and (F) DAPK1 is increased at 1 dpi in *SRR^fl/fl^* mice and not until 3 dpi in *TMEM119^creERT2^:SRR^fl/fl^* mice. (H,J) The ratio of phosphorylated to non-phosphorylated CREB is significantly upregulated in *TMEM119^creERT2^:SRR^fl/fl^* but not *SRR^fl/fl^* male mice after injury. (I,J) BDNF is significantly increased at 1 and 3 dpi in *TMEM119^creERT2^:SRR^fl/fl^*male mice, prior to upregulation in *SRR^fl/fl^* mice at 3 dpi. (K) GluN2B is downregulated in isolated hippocampal membrane fractions in male and female WT mice at 3 dpi, however (L) the ratio of phosphorylated GluN2B is increased in male but decreased in female mice at 3 dpi. *p<0.05, **p<0.01, ***p<0.001. Two-way ANOVA with Tukey’s multiple comparison test; n=4-5. Values represent mean ± SEM

Given the previously observed sex differences, we next examined NMDAR subunit levels in isolated plasma membrane fractions from hippocampal tissues of WT male and female mice. Interestingly, there was a significant downregulation of GluN2A (Figure S6F) and GluN2B (Figure 2K) in both male and female mice at 3 dpi, which could indicate self-regulatory internalization from the plasma membrane or loss from damaged spines. However, the ratio of pGluN2B to GluN2B was significantly increased in male mice but decreased in female mice at 3 dpi (Figure 2L), indicating enhanced activity of the remaining GluN2B containing NMDARs in male but not female mice.

To further examine the role of the GluN2B subunit in male mice after injury, we employed *CamKII^creERT2^:Grin2b^fl/fl^*mice, where tamoxifen treatment resulted in a 50-75% reduction of GluN2B levels in whole hippocampal lysate relative to Grin2b^fl/fl^ controls (Figure 3A). GluN1 and GluN2A levels remained unchanged, confirming a subunit-specific knockout (Figure S7 and S8A,B). There was a ∼50% reduction in GluN2B in the plasma membrane of *CamKII^creERT2^:Grin2b^fl/fl^*sham mice compared to *Grin2b^fl/fl^* sham mice (Figure 3B; Figure S7), where Na+/K+ ATPase expression was 2-4-fold enriched relative to whole lysates, validating successful membrane fraction isolation (Figure S8D).

**Figure 3.**
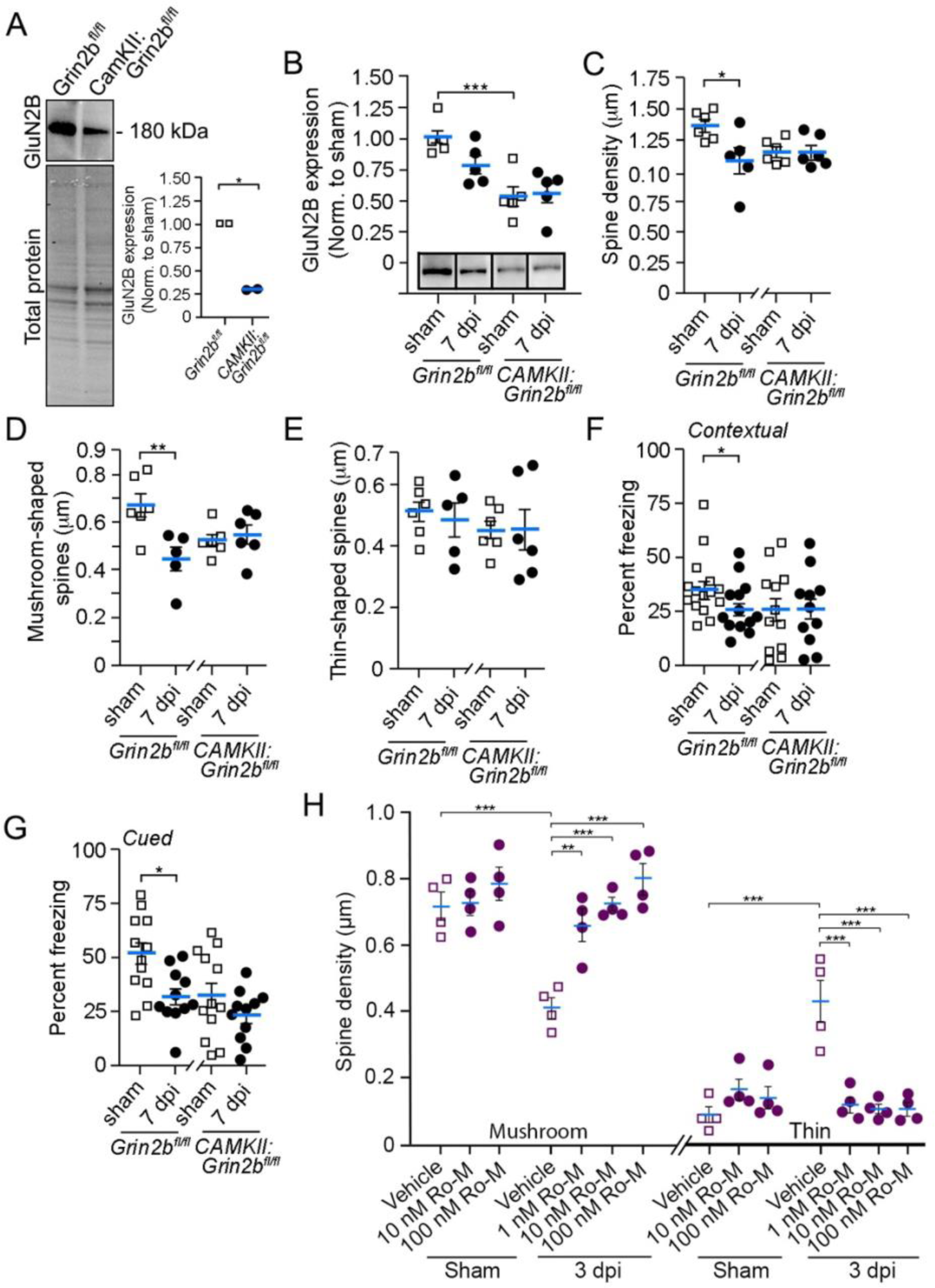
Inhibition of the GluN2B NMDAR subunit rescues synaptic damage and cognitive deficits after CCI injury. (A) Western blot analysis of whole hippocampal lysates shows reduced GluN2B expression in *CamKII^creERT2^:Grin2b^fl/fl^* mice. (B) Western blot analysis showing reduced GluN2B expression in isolated plasma membrane lysate from *CamKII^creERT2^:Grin2b^fl/fl^*mice compared to *Grin2b^fl/fl^* mice. (C) Reduced spine density in *Grin2b^fl/fl^* but not *CamKII^creERT2^:Grin2b^fl/fl^*male mice resulting from loss in (D) mushroom-shaped spines at 7 dpi but not (E) thin-shaped spines. (F) Contextual and (G) cued freezing are reduced in *Grin2b^fl/fl^* but not *CamKII^creERT2^:Grin2b^fl/fl^*male mice at 7 dpi. (H) Ro-maleate treatment leads to a rescue of mushroom- and thin-shaped spines at 3 dpi compared to vehicle treatment. *p<0.05, **p<0.01, ***p<0.001. Two-way ANOVA with Tukey’s multiple comparison test; (A) n=4, (C-E) n=5-6, (I) n=5. (B, F-G) Student’s t-test; (B) n=2, (F-G) n=10-15. Values represent mean ± SEM.

Control *Grin2b^fl/fl^* mice showed a ∼25% reduction in hippocampal CA1 dendritic spine density at 7 dpi (Figure 3C). This is due to the loss of mature mushroom-shaped spines that are preserved in *CamKII^creERT2^:Grin2b^fl/fl^* mice (Figure 3D). Thin-shaped spines were unchanged in both genotypes (Figure 3E), likely because they were already eliminated by the 7-dpi time-point. At a functional level, *Grin2b^fl/fl^* mice showed CCI-induced deficits on contextual (Figure 3F) and cued (Figure 3G) fear conditioning behavior at 7 dpi. *CamKII^creERT2^:Grin2b^fl/fl^* showed no injury-induced impairments in learning and memory; however, they exhibit reduced freezing in sham conditions relative to *Grin2b^fl/fl^* controls. This could be due to baseline differences in learning and memory resulting from pre-injury NMDAR alterations. To further support our hypothesis that GluN2B ablation has a therapeutic effect in CCI, we used a pharmacological approach with Ro-maleate 25-6981 (Ro), a GluN2B specific inhibitor. Vehicle saline or 1, 10, and 100 nM Ro were infused via Alzet osmotic minipumps for 3 dpi into the lateral ventricle of WT male mice. Golgi analysis showed significant losses in mushroom-shaped spines, corresponding to increases in thin-shaped spines in 3 dpi vehicle-treated mice (Figure 3H). Dendritic spine morphology was rescued by all three concentrations of Ro, supporting findings from *CamKII^creERT2^:Grin2b^fl/fl^* mice and the protective role of GluN2B inhibition after CCI injury.

### D-serine mediates synaptic pruning by regulating phosphatidylserine (PS), complement C1q, and the microglia GPR56 receptor levels after injury

Next, we examined the mechanisms downstream of NMDAR signaling that could lead to tagging of damaged synapses for elimination. Phosphatidylserine (PS) is a phospholipid localized to the inner leaflet of the plasma membrane during homeostasis. However, during apoptosis and in local synaptic compartments undergoing apoptotic-like mechanisms, PS is externalized to the outer plasma membrane to serve as an “eat-me” signal for phagocytes^36–40^. To assess whether this mechanism is affected by D-serine in CCI injury, we used a fluorescent dye, PSVue-480, which does not cross the plasma membrane and therefore binds externalized PS^41^. We delivered PSVue to the contralateral ventricle for 3 days using Alzet osmotic mini-pumps, where fluorescence was present at the injection site in sham mice and throughout the injured cortex at 3 dpi (Figure 4A, B). High magnification images show few PS+ puncta in the *stratum lacunosum moleculare (SLM*; Figure 4C-F) and *stratum radiatum* (*SR*; Figure S10M-P) of the CA1 hippocampus under sham conditions. At 3 dpi, there was an increase in PS+ puncta in the *SLM* of *SRR^fl/fl^* (Figure 4G-L) but not *Tmem119^creERT2^:SRR^fl/fl^* mice (Figure 4M-R), which is closely associated with the pre-synaptic marker Vglut1 and post-synaptic marker PSD-95. Quantification showed a significant increase in the percentage of Vglut1+/PSD95+ synapses that colocalized with PS at 3 dpi in control mice (Figure 4EE). This is 3.5-fold higher than that in *Tmem119^creERT2^:SRR^fl/fl^*mice, suggesting that microglial D-serine mediates PS externalization after *SLM* injury. Although similar trends were observed, no significant differences were observed in the *SR* (Figure 4FF; Figure S10 Q-U).

**Figure 4.**
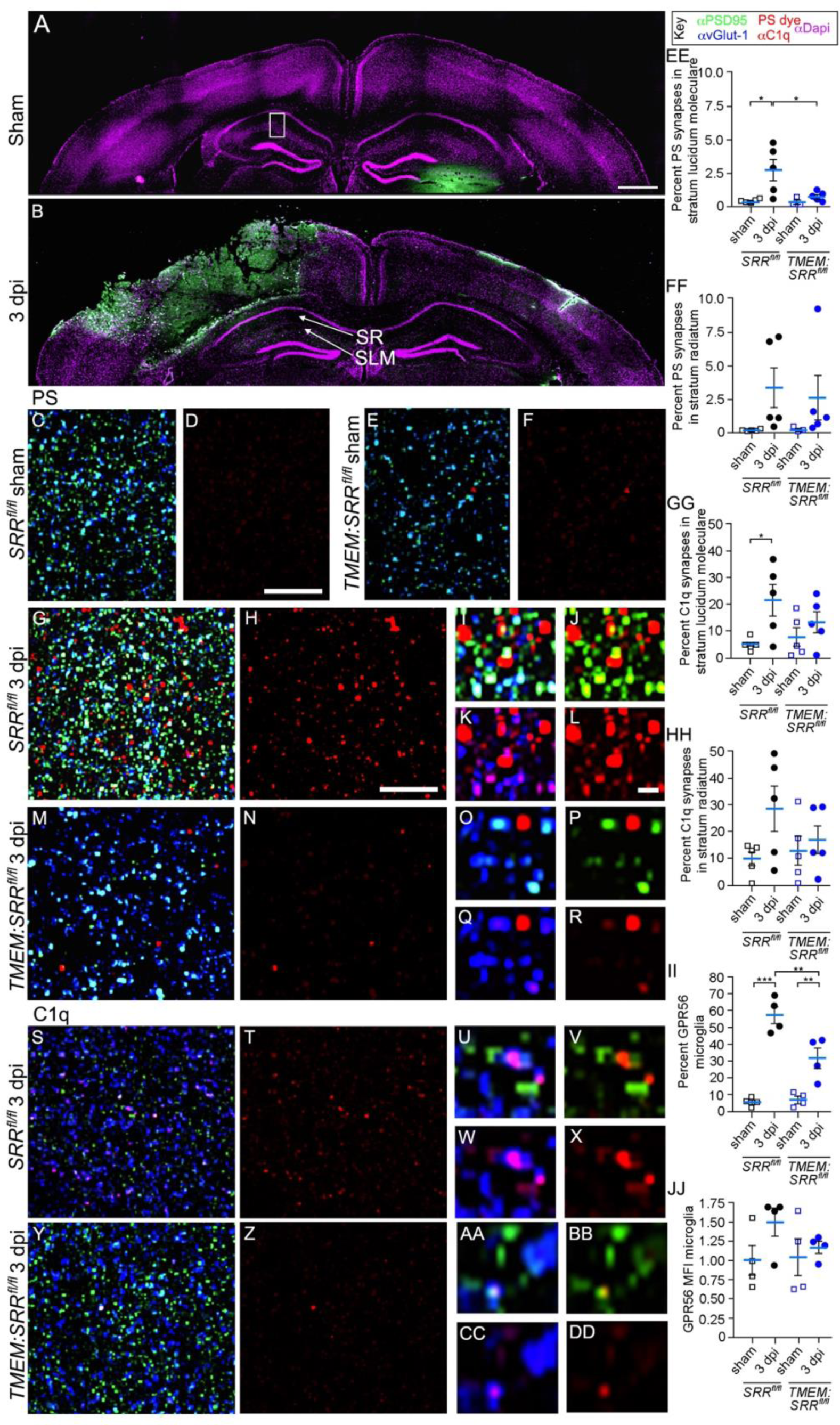
Upregulation of synaptic pruning components phosphatidylserine (PS), complement C1q, and GPR56 is mediated by microglial D-serine in the hippocampus of male mice. PSVue dye binds externalized PS following 3-day administration to the lateral ventricle in (A) sham or (B) CCI-injured mice. (C,E,G,I-L,M,O-R,S,U-X,Y,AA-DD) Anti-Vglut1 and Anti-PSD95 were used to identify the pre- and post-synaptic membranes. (C-F) High-magnification (63X) images show minimal labelling of externalized PS in the CA1 hippocampal *stratum lucidum moleculare* (*SLM*) of sham injured *SRR^fl/fl^* or *Tmem119^creERT2^:SRR^fl/fl^* mice, while increased PS was observed in (G-L) *SRR^fl/fl^* but not (M-R) *Tmem119^creERT2^:SRR^fl/fl^* mice. (S-DD) Anti-C1q expression was similar to PS expression. (EE-HH) Quantification measurements of PS and C1Q in the *SLM* and *SR*. (II, JJ) Hippocampal FACS microglia show elevated numbers and MFI of GPR56 expressing cells at 3 dpi in *SRR^fl/fl^* but to a lesser extent in *Tmem119^creERT2^:SRR^fl/fl^* mice. (Scale bar = (A) 500 μM, (D,H) 5 μM, (L) 1 μM. *p<0.05, **p<0.01, ***p<0.001. Two-way ANOVA with Tukey’s multiple comparison test; (EE-HH) n=3-5 mice/group each with 3 sections/area, (II,JJ) n=4. Values represent mean ± SEM.

Externalized PS interacts with C1q, a component of the complement system that is involved in synaptic pruning^41–43^. Here, we performed IHC staining of C1q with synaptic markers Vglut1 and PSD95 in the hippocampus, where C1q+ puncta increased in *SRR^fl/fl^* mice at 3 dpi relative to sham control mice in the *SLM* (Figure 4S-X; Figure S10A,B). Notably, fewer C1q+ puncta were visible in *Tmem119^creERT2^:SRR^fl/fl^*microglial mice at 3 dpi, comparable to the sham levels (Figure 4Y-DD; Figure S10C, D). Quantification showed a similar pattern as PS, where there was a significant 15-20% increase in C1q+ synapses at 3 dpi in the *SLM SRR^fl/fl^* mice but not *Tmem119^creERT2^:SRR^fl/fl^*mice (Figure 4GG). Combined analysis of the *SLM* and *SR* showed increased C1q colocalization with synapses after injury (Figure S10V); however, the *SR* alone showed trends but no significant changes (Figure 4HH; Figure S10E-L). Finally, we used FACS to quantify the expression of GPR56, a receptor for PS that plays a role in synaptic pruning^44,45^ (Figure S9). There was an increase in the percentage of microglia cells that expressed GPR56 after injury in both genotypes; however, significantly more microglia (∼25%) expressed GPR56 in *SRR^fl/fl^* than *Tmem119^creERT2^:SRR^fl/fl^* mice (Figure 4II). In addition, the MFI per GPR56+ microglia was 1.5-fold higher after injury in *SRR^fl/fl^* mice, although the difference was not statistically significant (Figure 4JJ). GPR56 expression was not observed in astrocytes (Figure S10W), suggesting that this is a microglia-specific mechanism of synaptic pruning after injury. Together, these findings demonstrate that the mechanisms for synaptic pruning, including PS, C1q, and GPR56, are upregulated after CCI injury and are modulated by D-serine.

### Single cell analysis of the CCI injured wild type male and female hippocampi

We performed single-cell RNA sequencing analysis on the hippocampi of male and female mice that underwent sham or CCI injury (3 dpi), using the Parse Bioscience Evercode Mega platform. Following quality control, the dataset consisted of 125,267 cells from sham tissue and 125,267 cells from 3 dpi tissues (Figure 5A). Following processing and clustering using Seurat v5, clear delineation was observed between 17 defined cell types and two undefined cell types (see color key), where CCI injury led to significant upregulation in both dividing microglia and infiltrating macrophages (Figure 5A,B; Figure S12A). Increased cell numbers after CCI injury were observed in both sexes, primarily due to a greater number of microglia and macrophages (Figure 5B). We next performed a subcluster analysis on the two key glia cell types in the tetrapartite synapse, microglia and astrocytes (Figure S12B,C). Unique astrocytic or microglial subclusters were not identified between male and female mice; however, differences were observed between the sham and CCI groups (Figure 5C). Specifically, we identified eight subclusters of astrocytes (A0-A7) and seven subclusters of microglia (M0-M6), where A2 and M1-M6 were all unique to CCI injury induction (Fig. 5C). We then determined whether there were unique and significant differences in the differential gene expression between male and female mice. Figure S13 shows differences in total upregulated or downregulated genes (open bars) as well as in genes unique to each sex (gray bars) observed in subclusters present in both sham and CCI conditions, where most genes were uniquely expressed. For subclusters uniquely observed only in CCI conditions, we compared the differences between the male and female CCI groups. Pathway Gene Ontology (GO) analysis was performed for both sham-CCI and CCI-CCI groups (Figure 5D-M; Figure S14; Table S1 with supplemental file 1).

**Figure 5.**
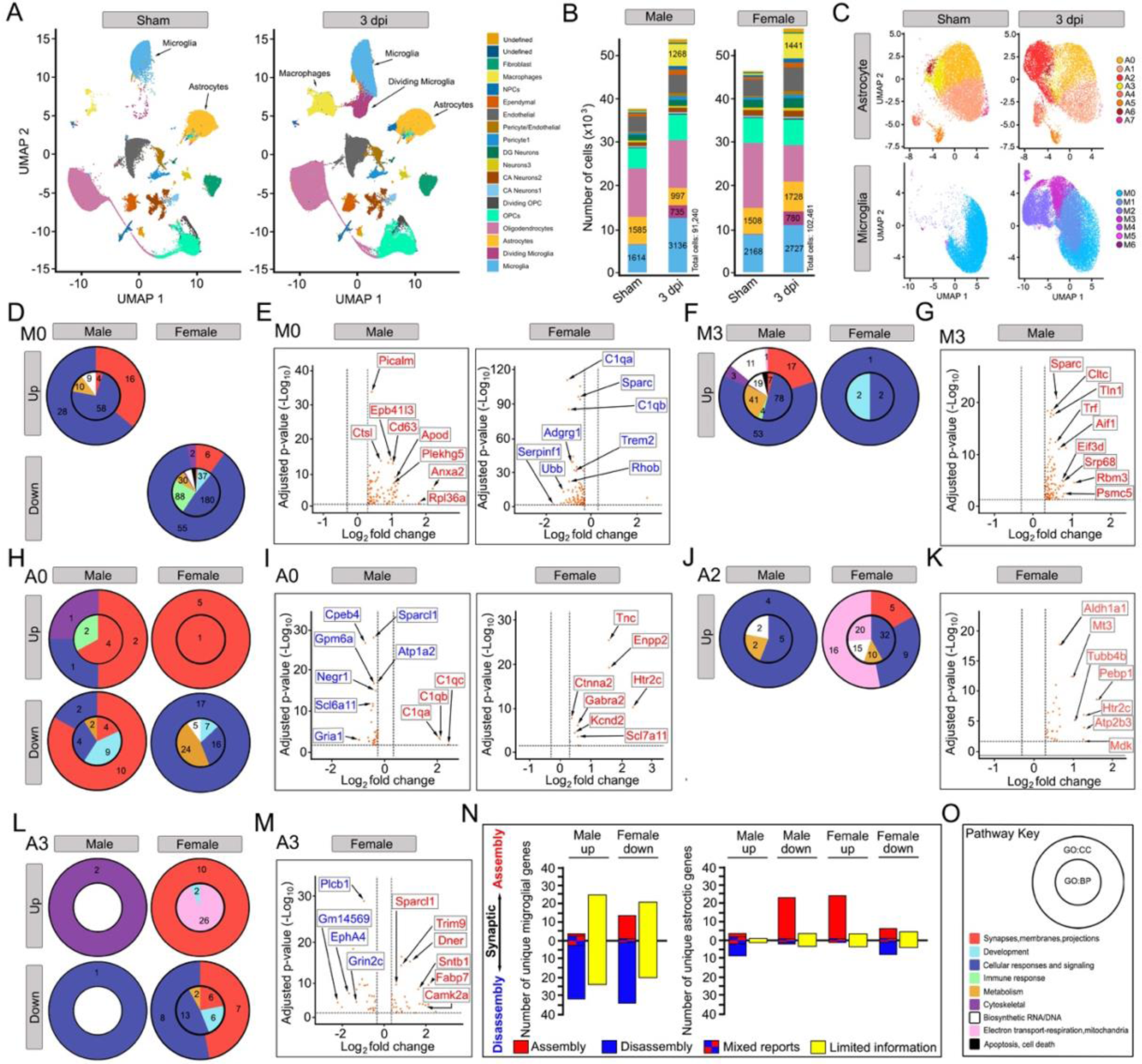
Single cell RNA-seq. analysis of male and female hippocampi. (A) UMAP cell distribution of sham and 3 dpi hippocampi shows 19 cell types. (B) Number of sham and 3 dpi cells in male and female mice shows increased cells after CCI injury. (C) UMAP of astrocyte and microglia subclusters show differences between sham and CCI injury groups. Upregulation and/or downregulation of synaptic-related pathways comparing sham to CCI injury were observed in male and/or female (D) microglial subclusters M0 and (H,L) astrocyte subclusters A0 and A3. Selective upregulated (red labels) genes and/or downregulated (blue labels) genes were observed in male and/or female (E) M0 and (G) M3 microglia and (I,K,M) A0 and A3 astrocytes. Subclusters (F,G) M3 and (J, K) A2 were not present in sham conditions, so comparisons are between CCI male and CCI female mice. (N) Summary of up- and down-regulated male and female genes from microglia or astrocyte populations, where red indicates genes associated with synaptic assembly, blue indicates synaptic disassembly, checkered blue/red indicates mixed functions, and yellow indicates synaptic-related genes with poorly defined synaptic functions. (O) Pathway key showing the cellular component (CC) in the outer pie and biological processes (BP) in the inner pie, where consolidated pathways are color-coded. Numbers in each pie region represents the number of consolidated GO pathways.

Specifically, we examined the biological processes (BP; inner ring) or cellular components (CC; outer ring) of the unique genes in males and females for each sub-cluster (Figure 5D,F,H,J,L; Figure S14B,E,F,G,J,K). Pie charts were developed based on consolidated GO terms (see key in Fig. 5O, Table S1 with supplemental file 1). Microglia subclusters showed unique differences in synaptic-related genes (red in pie; numbers represent the number of nonconsolidated GO pathways; Table S1 with supplemental file 1). Unique synaptic-related genes (orange dots) are graphed in corresponding figures to demonstrate the numbers of up- (red labeled) and downregulated (blue labeled) genes for each subcluster and sex based on ±0.3 Log2 fold change and <0.05 adjusted P-value (Figure 5E,G,I,K,M). The gene labels only represent selected genes with a greater adjusted P-value or Log2 fold change, where the entire list of significantly altered genes is found in supplemental file 1. Synaptic-related genes in microglia were primarily localized to M0, M2, and M3 subclusters, where uniquely upregulated synaptic genes were observed in male but not in female mice (Figure 5D-G; Figure S14K; Table S2). It should be noted that M0 was compared to sham controls, while M2 and M3 were direct comparisons between male and female CCI conditions. Interestingly, female M0 microglia downregulated synaptic-related genes, including the pruning genes complement genes C1q and Trem2 (Table S2; 5D, E).

In astrocyte subclusters A0, A1, A3, A5, and A6, we also observed pathways that were uniquely up- or downregulated in males versus females, and vice versa (Figure 5H-M; Figure S14A-G). A0 and A1 showed increases in a small pro-pruning gene set, mainly C1q, which was absent in female astrocytes (Figure 5H,I; Figure S14B). Conversely, synaptic-related genes in female A0, A2, and A3 astrocytes were generally pro-synaptic (Figure 5H-M). Since subcluster A2 was not present in sham conditions, the upregulation of synaptic-related genes in females was compared with that in male CCI mice (Figure 5J, K).

To summarize these concepts, we examined whether genes differentially expressed by microglia or astrocytes preferentially target synaptic assembly or disassembly based on a literature review (Figure 5N; Table S2 with supplemental file 1). In microglia, a greater number of uniquely upregulated genes in males and downregulated genes in females tended towards synaptic disassembly. However, many unique genes with synaptic associations have limited functional information (yellow bar), and it is difficult to determine the role of this group of genes without further mechanistic studies. For example, one large group of genes with limited information unique to males was the upregulation of 23 ribosomal genes, which could indicate greater translational responses after injury (Table S2 with supplemental file 1). Genes uniquely expressed in male astrocytes showed tendencies towards synaptic dysfunction, as the number of genes that supported synaptic disassembly (blue bars) was preferentially upregulated, whereas synaptic assembly (red bars) was downregulated (Figure 5N). Genes uniquely expressed in females showed opposite tendencies. In short, we recognize that a greater understanding of gene function between males and females is warranted. Importantly, our single-cell data strongly support biological functional studies demonstrating neuroprotection in females after injury and provide a better understanding of the genes that may underlie these mechanisms.

### Single cell analysis of the CCI injured microglial D-serine deficient hippocampi

To examine the effects of microglial D-serine on CCI injury in male mice, we performed scRNA-seq analysis on the hippocampi of *Tmem119^creERT2^:SRR^fl/fl^* mice as compared to *SRR^fl/fl^*control mice that underwent sham and CCI injury (3-dpi) using the Parse Bioscience Evercode Mega platform. We observed a similar distribution in cell number (Figure 6A) and cell percentage (Figure S15A) in sham and 3 dpi cell types between *SRR^fl/fl^* (56,820 total cells) and *Tmem119^creERT2^:SRR^fl/fl^* (48,733 total cells) mice. Similar to WT mice, an increase in cell number was observed in both groups after CCI injury. Differential expression was calculated for microglia and astrocytes between *SRR^fl/fl^* to *Tmem119^creERT2^:SRR^fl/fl^* (Figure 6B-D; supplemental file 1). We observed an increase in synaptic-related genes (red labels show representative genes), including *C1q*, associated with the presence of D-serine in both dividing and non-dividing microglia (i.e. *SRR^fl/fl^* mice). This suggests that microglial D-serine deficiency reduces synaptic-related gene expression. Interestingly, astrocytes also showed increases in synaptic-related genes in *SRR^fl/fl^*compared to *Tmem119^creERT2^:SRR^fl/fl^* mice, supporting the possibility that microglial D-serine release alters astrocytic gene function (Figure 6D).

**Figure 6.**
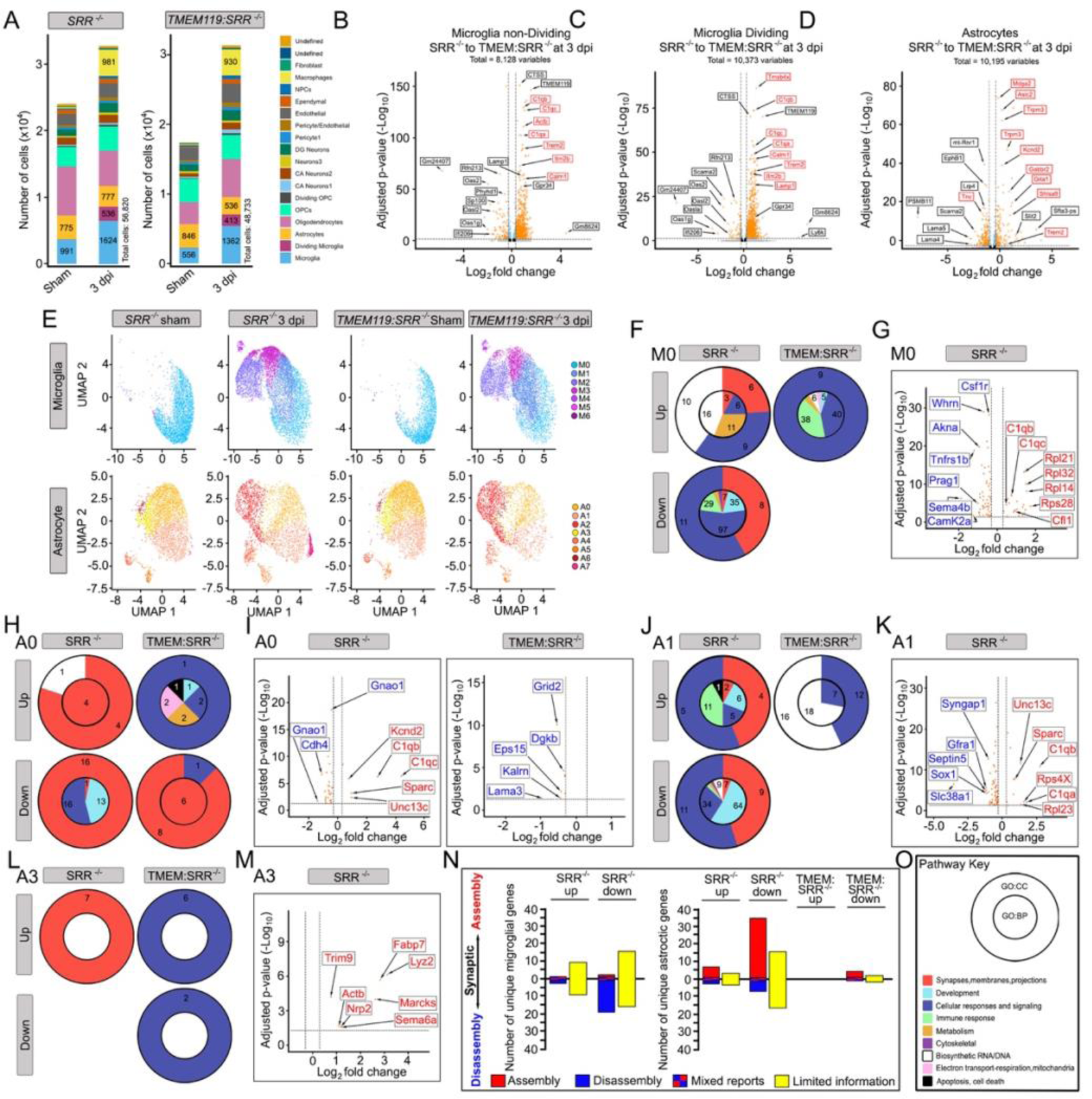
Single cell RNA-seq. analysis of *SRR^fl/fl^* and *Tmem119^creErt2^:SRR^fl/fl^* hippocampi. (A) Numbers of sham and 3 dpi cells in *SRR^fl/fl^* and *Tmem119^creErt2^:SRR^fl/fl^*mice shows increased numbers after CCI injury. Volcano map of (B) non-dividing microglia, (C) dividing microglia, and (D) astrocytes where +Log_2_ fold-change represents select genes upregulated in *SRR^fl/fl^* mice (red labels represent synaptic-related genes) and -Log_2_ fold-change represents genes upregulated in *Tmem119^creErt2^:SRR^fl/fl^*mice. (E) UMAP of astrocyte and microglia subclusters show little differences between sham and CCI injured *SRR^fl/fl^* and *Tmem119^creErt2^:SRR^fl/f^* groups. Upregulation and/or downregulation of synaptic-related pathways comparing sham to CCI injury were observed in *SRR^fl/fl^* and *Tmem119^creErt2^:SRR^fl/f^* (F) microglial subclusters M0 and (H,J,L) astrocyte subclusters A0, A1, and A3. Selective upregulated (red labels) genes and/or downregulated (blue labels) genes were observed in *SRR^fl/fl^* and *Tmem119^creErt2^:SRR^fl/f^*(G) M0 microglia and (I,K,M) A0, A1, and A3 astrocytes. (N) Summary of upregulated and downregulated genes associated with *SRR^fl/fl^* and *Tmem119^creErt2^:SRR^fl/fl^*mice from microglia or astrocyte populations, where red indicates genes associated with synaptic assembly, blue indicates synaptic disassembly, checkered blue/red indicates mixed functions, and yellow indicates synaptic-related genes with poorly defined synaptic functions. (O) Pathway key showing the cellular component (CC) in the outer pie and biological processes (BP) in the inner pie, where consolidated pathways are color-coded. Numbers in each pie region represents the number of consolidated GO pathways.

Subcluster analysis was performed on both microglia and astrocytes (Figure S12B,C), where subcluster types and distributions were similar to those in WT mice (Figure 5C, 6E). *SRR^fl/fl^* and *Tmem119^creERT2^:SRR^fl/fl^* mice were similar, except for A7, which was present in CCI-injured *SRR^fl/fl^* mice but not CCI injured *Tmem119^creERT2^:SRR^fl/fl^* mice (Figure 6E). Pathway analysis (Key in Figure 6O; Table S3) demonstrated that CCI injury altered synaptic-related genes in control *SRR^fl/fl^* mice, but not in the absence of microglial D-serine (Figure 6G; Figure S16A). In particular, *C1q* and many ribosomal genes were uniquely upregulated in microglia from *SRR^fl/fl^* but not *Tmem119^creERT2^:SRR^fl/fl^*mice. Male mice from both genotypes showed a similar increase in ribosomal genes, supporting the importance of these shared regulatory genes in synaptic damage. Pro-pruning *C1q* genes were upregulated in many microglia subclusters (M0, M1, M2, M3, and M6), while the phagocytic receptor *Trem2* was specific to M1 (Figure 6A-D; Table S4 with supplemental file 1). Interestingly, many astrocytic sub-clusters also demonstrated synaptic-related alterations, where A0, A1, and A3 showed changes in synaptic-related genes in injured *SRR^fl/fl^* mice (Figure 6H-M; Figure S16H,I). *Tmem119^creERT2^:SRR^fl/fl^* mice showed downregulation of nine synaptic genes in cluster A0; however, the 98 genes up- or down-regulated in *SRR^fl/fl^* mice were not significantly changed in the absence of microglial D-serine (Figure 6H-M; Table S4 with supplemental file 1).

Analysis of all microglial genes showed no significant changes in *Tmem119^creERT2^:SRR^fl/fl^* mice, whereas *SRR^fl/fl^* mice showed both up- and down-regulated genes, including a large number of synaptic genes with limited functional information (yellow bar) (Figure 6N; Table S4 with supplemental file 1). Similar changes were observed in astrocytes, where few genes were significantly altered in *Tmem119^creERT2^:SRR^fl/fl^* mice. In short, our study supports the role of D-serine in regulating synaptic-regulated genes in glia that are associated with tetrapartite synapses and synaptic disassembly (Figure 6N).

### Genetic and biochemical alterations in human TBI patients support factors that regulate synaptic losses and pruning

Finally, to begin to translate our findings into TBI patients, we examined bulk RNA-sequencing transcriptional changes in perilesional tissues resected from 32 neurotrauma patients and 3 human cortical control samples (Figure 7A; Table S5). As anticipated, differential expression analysis revealed transcriptomic changes after TBI; however, we also observed differences within our tested TBI population. Particularly, analyzing samples based on the time of hospital admittance to surgery as <1 day or >1 day revealed differences in the gene profile (Figure 7B-D).

**Figure 7.**
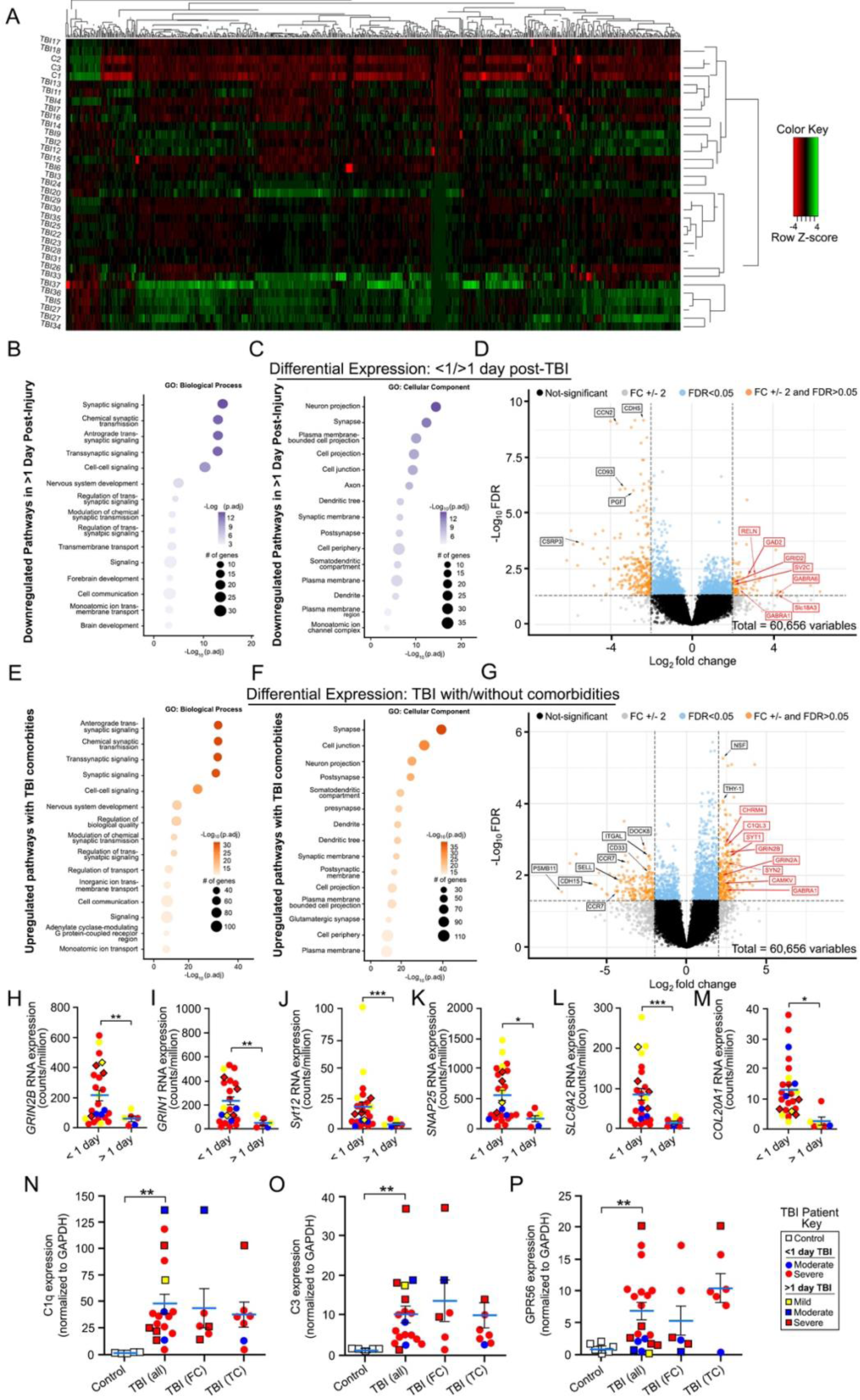
Human TBI patients have altered expression of synapses and synaptic pruning components. (A) Heatmap of differentially expressed genes from bulk RNA sequencing of 32 cortical TBI tissue samples and 3 controls. (B,C) Pathway analysis of differentially expressed genes between samples from patients that underwent surgery in <1 day after hospital admittance versus patients that underwent surgery >1 day after admittance. (B) Pathways from GO:biological process and (C) GO:cellular component both suggest lower expression of genes related to synaptic pathways in samples from later timepoints after injury. (D) Volcano plot depicting differentially expressed genes between samples acquired less than a day versus more than a day after TBI, where genes with a positive fold change (FC) are enriched in less than a day samples. (E,F) Pathway analysis of differentially expressed genes between samples from patients with and without known CNS comorbidities. (E) Pathways from GO:biological process and (F) GO:cellular component both suggest higher expression of genes related to synaptic pathways in patients with CNS comorbidities. (G) Volcano plot depicting differentially expressed genes between samples with and without known CNS comorbidities. (H-M) Distribution pattern of select synaptic related genes that are downregulated in patients that underwent surgery >1 day after hospital admittance relative to patients that underwent surgery in < 1day. Western blot analysis shows upregulated protein levels of synaptic pruning components (N) C1q, (O) C3, and (P) GPR56 in TBI samples relative to controls. *p<0.05, **p<0.01, ***p<0.001. Mann-Whitney test; (H-M) n = 6-25, N-O n= 3-19. Values represent mean ± SEM.

We observed that many significantly upregulated genes in synaptic signaling, synaptic membrane interactions, and transmission occurred in TBI patients that acutely underwent surgery (i.e., <1 day) as compared to patients who underwent surgery after the first day. Conversely, more chronic surgical patients showed an increase in the expression of inflammatory genes (Figure 7 D). We also observed a difference within the <1 day to surgery group, where TBI patients with previous or existing CNS comorbidities (such as depression, bipolar disorder, Parkinson’s disease, substance abuse, PTSD, schizophrenia, anxiety, ADHD, frontotemporal dementia, and epilepsy; Table S5) also showed a differential increase in synaptic-related genes compared with patients without reported comorbidities (Figure 7E-G). The volcano plots show representative synaptic genes in red (Figure 7D, G), and the distribution of six representative genes, namely *GRIN2B, GRIN1, SYT12, SNAP25, SLCl8A2, and COL20A1,* is shown for mild, moderate, and severe TBI (Figure 7H-M). Color coding depicts TBI severity based on the Glasgow Coma Scale (GCS), where yellow indicates mild TBI (13-15 GCS), blue indicates moderate TBI (9-12 GCS), and red indicates severe TBI (3-8 GCS). Triangles represent female patients, whereas circles indicate male patients. There was no observable correlation between GCS severity and synaptic-related gene expression.

To assess the protein levels of synaptic pruning components, we performed Western blot analysis on TBI versus control samples (Figure 7N-P). Control samples included a whole brain sample (C1) used to normalize across gels, as well as 5-6 quantified cortical samples (C2-C7) (Table S5). Analysis of C1q protein levels indicated a significant 48-fold increase in TBI patient samples compared to controls, with a distribution ranging between 4.4-136-fold across patients (Figure 7N). We examined the expression of a second complement component, C3, which is involved in the tagging of damaged synapses downstream of C1q^42^, and found a significant 10-fold increase compared to the controls (Figure 7O). Finally, we found that GPR56 expression was significantly increased, on average, ∼6.9-fold in TBI samples, ranging from ∼0-20-fold (Figure 7P). Notably, GPR56 expression was the highest in samples from severely injured patients who underwent surgery in <1 day, suggesting that GPR56 is acutely activated after injury. Together, these results show that nearly all tested TBI patients had increased protein levels of C1q, C3, and GPR56, supporting our CCI model and the possibility that similar mechanisms of synaptic pruning are conserved across mouse and human TBI.

## DISCUSSION

Synaptic damage is a hallmark of TBI pathology and can lead to persistent impairments in cognitive function^3–7^. Here, we demonstrate that microglial D-serine alters the expression of GluN2B NMDAR signaling pathways after injury, likely leading to the overactivation of excitatory neurotransmission and subsequent tagging of spines for elimination. Upregulation of the GluN2B subunit has been implicated in synaptic dysfunction in many neurodegenerative models^26–28^ and excitotoxic cell death in TBI pathologies^46–49^. Our findings also demonstrate that cognitive dysfunction results from diffuse, widespread synaptic loss and not the focal loss of cortical tissues after brain impact injury. In fact, targeted synaptic elimination is independent of neuronal death or inflammatory responses but is dependent on the quadpartite synaptic microenvironment where activated astrocytes and microglia are involved. The implications of these findings may extend beyond severe TBI to implicate cognitive dysfunction associated with other diseases and disorders in which activated gliosis is present.

Glutamate excitotoxicity is an expected mechanism of acute neuronal cell death through the hyperactivation of NMDARs, and broad-spectrum NMDAR inhibitors can reduce cell death in animal injury models^50–52^. However, these studies failed in clinical trials, which is believed to result mainly from the acute timing of glutamatergic excitotoxicity minutes after the initial trauma^53,54^. As broad inhibition of NMDARs can also exacerbate damage due to their role in pro-survival signaling, specific GluN2B inhibition may be more promising, especially at lower concentrations aimed at preventing ongoing synaptic damage beyond the initial impact^34,53,55,56^. While glutamate levels stabilize after brain injury, we show prolonged increases in the NMDAR co-agonist D-serine^6,7^. The ability of glial D-serine to modulate NMDAR expression and downstream signaling pathways has important implications in many neurological conditions, as NMDAR dysfunction is involved in many neurodegenerative conditions beyond TBI^57^. This can provide a novel therapeutic target upstream of NMDARs, where inhibiting glial D-serine release enables the modulation of NMDAR activity without affecting excitatory neurotransmission.

The complement system, as well as phosphatidylserine (PS), has been studied for its role in circuitry refinement during development, synaptic plasticity in adulthood, and synaptic loss in neurodegeneration and injury^37,41,42,44,58–67^. However, a clear activator of these signals has not been previously demonstrated. Our studies suggest that glial D-serine initiates a targeted mechanism of synaptic elimination following brain trauma rather than broad synaptic loss from general inflammatory damage. The link between GluN2B and PS externalization is likely due to excessive calcium influx, which leads to the cleavage of flippases and scramblases that regulate PS localization, possibly in a caspase-dependent manner^36–40^. Once PS is externalized, it can bind complement components such as C1q or C3, which we demonstrate are upregulated and released by microglia and astrocytes after TBI^41,43,68,69^. The PS-complement complex has been shown to interact with receptors such as GPR56^43–45,70^, a microglial receptor that is associated with phagocytosis. Our findings indicate that D-serine may contribute to a positive feedback loop in synaptic pruning, as microglial SRR-ablated mice had downregulated GPR56 expression after injury. This suggests that microglia actively tailor their responses to match the levels of synaptic damage, where higher expression of the ligand PS likely induces the expression of its receptor. The identification of microglial GPR56 as a downstream response to D-serine provides an interesting avenue for future studies, where functional effects remain to be established. It is also likely that other microglial receptors, including Trem2 and C3R^41,63^, which have upregulated RNA expression after injury, are involved in the pruning process. The degree to which these receptors and their ligands have overlapping functions after TBI remains unclear. Therefore, it is important to evaluate whether the elimination of one receptor, GPR56, leads to compensatory activity by other microglial receptors. Furthermore, it cannot be overlooked that, in addition to microglia, astrocytes play an active role in synaptic pruning^71^, and further experiments are warranted to determine the interactions between both cell types after injury.

We did not observe D-serine-mediated synaptic damage in female mice, where glial SRR, the initiator of synaptic pruning, was differentially reduced primarily in microglia and, to a lesser extent, astrocytes. This may be because lower levels of glial D-serine reduced the total extracellular levels of D-serine below the threshold amount necessary to induce synaptic damage, which in turn resulted in reduced GluN2B phosphorylation, preserved dendritic morphology, and improved cognitive outcomes after TBI. Single-cell analysis between male and female hippocampi show that both microglia and astrocytes have unique sex-specific gene expression patterns, where CCI injury leads to increased numbers of synaptic disassembly genes in male astrocytes and microglia, while males also have reduced numbers of assembly genes in astrocytes. Female astrocytes and microglia show the opposite effect in synaptic gene responses with expression of unique genes not observed in males. When comparing genes associated with assembly or disassembly between male/female mice and *SRR^fl/fl^/Tmem119^creERT2^:SRR^fl/f^*mice, we do observe a number of conserved unique gene variances; however, the majority of genes associated with protection in female astrocytes or microglia are not identical to *Tmem119^creERT2^:SRR^fl/fl^*male mice. One major exception is the expression of *C1q* genes. It became quite evident that expression of these pro-pruning components corresponded to our mechanistic findings, where C1q is upregulated in astrocytes and microglia of male and *SRR^fl/fl^* mice, but reduced in female and *Tmem119^creERT2^:SRR^fl/fl^* mice. Importantly, in human TBI tissues, all analyzed patients showed increased complement levels as compared to control human cortical tissues.

Neuroprotective responses in females have been observed in a number of studies where reduced microglial reactivity is associated with female pathologies, and anti-inflammatory responses are associated with estrogen and progesterone levels ^16,23–25^. Our studies support this phenomenon after TBI and demonstrate unique glial gene expression variances that are associated with synaptic functions. This raises the possibility that female circulating hormones contribute to the reduced production of microglial D-serine after injury, although further studies are needed.

Moreover, we have previously shown that SRR and Slc1a4 are upregulated in human TBI^7^. Here, we identify similar increases in synaptic pruning components. In particular, C1q and C3 are increased in most TBI tissues, similar to that observed in our mouse model of TBI. The involvement of complements in tagging cells or cell membranes for phagocytosis^72^ or synaptic elimination have been previously described^68^. However, given the multifaceted role of the complement system after injury, it is difficult to demonstrate that C1q and C3 are directly involved in synaptic tagging and elimination after TBI^68^. Our observation that GPR56 is upregulated in mouse and human tissues, together with the association of PS and complement C1q with pre- and post-synaptic membranes, supports the concept that these factors are involved in TBI-induced synaptic pruning. Analysis of transcriptomic alterations supports acute alterations in synaptic genes, including *GRIN2B*, which supports the translation of our mouse findings to human patients with TBI. In addition, there is global downregulation of synaptic genes in patients with TBI over time. This is demonstrated by our comparison of patients in whom tissues were removed from the perilesional cortex in acute versus more chronic periods. This finding also supports the progressive loss of synaptic integrity over time, similar to that observed in our mouse model. We cannot exclude the possibility that synaptic genes are affected by neuronal loss in acute TBI patients; however, we observed comparable differences in cell death genes between acute and chronic TBI patients. Thus, it is possible that the treatment of human TBI patients with glial D-serine release inhibitors, such as L-4-Chlorophenylglycine, could be protective if administered in acute periods after brain injury. It is also possible that D-serine functions as an important biomarker after injury to determine the extent of cognitive dysfunction; however, extensive biomarker analysis is needed in both mice and humans.

In conclusion, we identified a novel mechanism by which microglial D-serine induces synaptic damage after injury. As reactive gliosis and NMDAR dysfunction are hallmarks of many neurodegenerative conditions, our findings have important implications for synaptic loss beyond TBI. Notably, we identified sex differences in TBI outcomes, which could suggest altered production or effects of microglial D-serine in females. Together, these findings will pave the way for the development of targeted treatments to mitigate synaptic dysfunction after TBI and to improve patient outcomes.

## Supporting information

All supplemental data

## SUPPLEMENTAL INFORMATION

Supplemental information can be found below.

## ACKNOWLEDGEMENTS

The authors thank Maria M. Quiala-Acosta, Jose Mier and Maria L. Cepero for technical assistance, animal husbandry, surgical procedures, as well as tissue harvesting and analysis. We would also like to thank Dr. Oliver Umland for expertise in conducting the flow cytometry and FACS studies. This work was supported by the Miami Project to Cure Paralysis, National Institute of Health/National Institute of Neurological Disorders and Stroke (NS098740; DJL), and the Lois Pope Life Fellowship.

## AUTHOR CONTRIBUTION

DA and DJL contributed to the overall study design, data collection, analysis, and/or interpretation. CAD and CMA designed, performed, and analyzed electrophysiology experiments. GFB processed samples for plasma membrane isolation. JRJ, BAM, and JGC contributed to the acquisition of human TBI tissues. OOF performed SRR immunofluorescence experiments. JBF and KPD conducted Multiplex analysis. AJG analyzed single cell and bulk RNA sequencing data. HW contributed to conceptual design and analysis. DA and DJL wrote the manuscript with assistance from all other authors.

## STAR + METHODS

### KEY RESOURCE TABLE

| REAGENT or RESOURCE | SOURCE | IDENTIFIER |
| --- | --- | --- |
| **Antibodies** |  |  |
| Rabbit anti-PSD-95 | ThermoFisher | 51-8900 |
| Guinea pig anti-Vglut1 | Synaptic Systems | 135304 |
| Rat anti-C1qa | Novus Biologicals | NB200-539 |
| Rabbit anti-BDNF | Abcam | Ab9793 |
| Rabbit anti-phospho-ERK1/2 | Cell Signaling | 9101S |
| Rabbit anti-ERK1/2 | Cell Signaling | 9102 |
| Rabbit anti-NR1 | Millipore | 05-432 |
| Rabbit anti-NMDAR1 | Millipore | Ab109182 |
| Rabbit anti-NR2A | Millipore | 07-632 |
| Rabbit anti-NR2B | Millipore | 06-600 |
| Rabbit anti-GRIN2B | ThermoFisher | 21920-1-AP |
| Rabbit anti-DAPK1 | Cell Signaling | 3008S |
| Rabbit anti-p-CREB | ABclonal | AP0019 |
| Rabbit anti-CREB | ABclonal | A11064 |
| Rabbit anti-Sodium Potassium ATPase | Abcam | AB76020 |
| Rabbit anti-Phospho-N2B,Ser1303 | Millipore | 07-398 |
| Rabbit anti-C1qA | Abcam | Ab189922 |
| Rabbit anti-C3 | Invitrogen | PA5-21349 |
| Mouse anti-GPR56 | Biolegend | 358205 |
| Mouse anti-SRR | BD Biosciences | 612053 |
| Mouse anti-SRR | Santa Cruz Biotech | Sc-365217 |
| Rabbit anti-Slc1a4 | Cell Signalling | 8442S |
| Mouse anti-GAPDH | Santa Cruz Biotech | 47724 |
| Goat anti-rabbit | Jackson Immuno | 111-035-003 |
| Goat anti-mouse | Jackson Immuno | 115-035-003 |
| Alexa Fluor™ 488 goat anti-mouse | ThermoFisher | A1101 |
| Alexa Fluor™ 647 goat anti-rabbit | ThermoFisher | 1524909 |
| Alexa Fluor™ 594 goat anti-guinea pig | ThermoFisher | 2540867 |
| Alexa Fluor™ 647 goat anti-rat | ThermoFisher | A21247 |
| CD45-PerCP-Vio 700, REAfinity™ | Miltenyi Biotech | 130-110-801 |
| CD11b-APC | Biolegend | 101212 |
| ACSA2-PE | Miltenyi Biotech | 130-110-801 |
| GPR56-PE/Cy7 | Biolegend | 358205 |
| **Biological samples** |  |  |
| Cortical protein lysates | Novus Biological | NB820-59186,<br>NB820-60574,<br>NB820-59193,<br>NB820-59198,<br>NB820-59177 |
| Cortical RNA | Takara Bio | 636570, 636571,<br>636564 |
| **Chemicals, peptides, and recombinant proteins** |  |  |
| Tamoxifen | Sigma Aldrich | T5648 |
| PSVue-480 | Molecular Targeting Technologies | P-1003 |
| Ro 25-6981 maleate | Bio-Techne Corporation | 15-941-R |
| Paraformaldehyde | ThermoFisher | A11313.36 |
| Sucrose | VWR | 0355 |
| Fluoro-Gel mounting media | Electron Microscopy Sciences | 17985-10 |
| AffiniPure Fab Fragment Goat αMouse IgG (H+L) | Jackson Immuno | 715-007-003 |
| Sudan Black | Sigma Aldrich | 199664 |
| Protease inhibitor | Sigma Aldrich | 04693159001 |
| Phosphatase inhibitor | Sigma Aldrich | 04906845001 |
| NuPAGE™ LDS Sample Buffer | Thermo Fisher | NP0007 |
| 2-Mercaptoethanol | Sigma Aldrich | M6250 |
| SuperSignal West Pico PLUS Chemiluminescent Substrate | ThermoFisher | 34578 |
| Hibernate A | ThermoFisher | NC0176976 |
| DAPI | Miltenyi | 130-116-141 |
| NucBlue Fixed Cell DAPI | ThermoFisher | R37606 |
| FcR blocking reagent TruStain FCX™ PLUS | Biolegend | 156603 |
| Actinomycin D | Sigma Aldrich | A1410 |
| Triptolide | Sigma Aldrich | T3652 |
| Anisomycin | Sigma Aldrich | A9789 |
| RNAasin Plus RNase Inhibitor | Promega | N2615 |
| 5% BSA Fraction V | ThermoFisher | 15260037 |
| **Critical commercial assays** |  |  |
| MILLIPLEX Mouse Cytokine/Chemokine Magnetic Bead Panel | Sigma-Aldrich | MCYTOMAG-70K-PMX |
| Mouse MMP Magnetic Bead Panel 3 | Sigma-Aldrich | MMMP3MAG-79K |
| FD Rapid GolgiStain Kit | FD NeuroTecnologies | PK401 |
| Plasma Membrane Protein Isolation and Cell Fractionation kit | Invent Biotechnologies | SM-005 |
| Direct-zol RNA Miniprep kit | Zymo Research | R2051 |
| Quick Start Bovine Serum Albumin (BSA) Standard kit | Bio-Rad | 5000206 |
| 4%–20% stain-free gradient gel | Bio-Rad | 5678094 |
| Trans-Blot Turbo RTA Transfer Kit | Bio-Rad | 1704270 |
| Adult Brain Dissociation Kit | Miltenyi Biotech | 130-107-677 |
| Evercode™ Cell Fixation v2 | Parse Biosciences | ECF2101 |
| **Experimental models: Organisms/strains** |  |  |
| C57BL/6J wildtype mice | Jackson Laboratory | 000664 |
| C57BL/6-*Tmem119 em2(EGFP)Gfng* /J | Jackson Laboratory | 031820 |
| C57BL/6J-*Grin2b<sup>em5Lutzy</sup>*/J | Jackson Laboratory | 032664 |
| *SRR<sup>fl/fl</sup>* | Bennyworth et al., 2012<sup>69</sup> | N/A |
| B6;129S6-Tg(Camk2a-cre/ERT2)1Aibs/J | Jackson Laboratory | 012362 |
| **Software and algorithms** |  |  |
| FreezeFrame 5 | Actimetrics | https://actimetrics.com/products/freezeframe/ |
| EthoVisionXT | Noldus | https://www.noldus.com/ethovision-xt |
| Reconstruct | Harris Lab, Center for Learning and Memory, University of Texas<sup>30</sup> | https://synapseweb.clm.utexas.edu/software-0 |
| ClearView-GPU Deconvolution | Oxford Instruments | https://imaris.oxinst.com/products/clearview-gpu-deconvolution |
| Imaris 10.0 | Oxford Instruments | https://imaris.oxinst.com/products/image-analysis-software/ |
| Image Lab | Bio-Rad | https://www.biorad.com/en-us/product/image-lab-software |
| CytExpert | Beckman Coulter | https://www.beckman.pt/flow-cytometry/research-flow-cytometers/cytoflex/software |
| TrimGalore | Github | https://github.com/FelixKrueger/TrimGalore |
| STAR v2.5.2 | Dobin et al., 2013<sup>73</sup> | https://github.com/alexdobin/STAR |
| ComBat-seq | Zhang et al., 2020<sup>74</sup> |  |
| DESeq2 | https://genomebiology.biomedcentral.com/articles/10.1186/s13059-014-0550-8 | https://bioconductor.org/packages/release/bioc/html/DESeq2.html |
| ParseBiosciences-Pipeline.1.1.2 | Parse Biosciences | https://support.parsebiosciences.com/hc/en-us/sections/27082017166484-Parse-Pipeline |
| Seurat v5 | Hao et al., 2024<sup>74</sup> | https://satijalab.org/seurat/ |
| GProfiler | Kolberg et al., 2023<sup>74</sup> | https://biit.cs.ut.ee/gprofiler/gost |
| Prism 9 | GraphPad | https://www.graphpad.com/features |
| pClamp 11.2 | Molecular Devices | https://www.moleculardevices.com/products/axon-patch-clamp-system |
| **Other** |  |  |
| Zeiss Axiovert 200M fluorescence microscope | Zeiss | SM100309-MG1PH |
| Keyence fluorescence microscope | Keyence | BZ-X810 |
| Dragonfly high speed confocal microscope | Andor | https://andor.oxinst.com/products/dragonfly-confocal-microscope-system?ppc_keyword=confocal+imaging+system&gad_source=1&gclid=Cj0KCQjw-5y1BhC-ARIsAAM_oKk12DWeb708Aa95HSkuyZuFLOGnCElvXV29lpPk6vF5psAy4xd0AroaAnTGEALw_wcB |
| ChemiDoc MP Imaging System | BioRad | Version 2.3.0.07 |
| ChemiDoc Touch Imaging System | BioRad | Version 1.2.0.12 |
| gentleMACS™ Octo Dissociator | Miltenyi Biotech | 130-096-427 |
| MoFlo Astrios Cell Sorter | Beckman Coulter | C09766 |
| Cytoflex S flow cytometer | Beckman Coulter | B25982 |
| Leica 1200 VT vibratome | Leica Biosystems | https://www.leica-biosystems.com/us/research/vibratomes/leica-vt1200/ |
| Multiclamp 700B amplifier | Molecular Devices | https://www.moleculardevices.com/products/axon-patch-clamp-system/amplifiers/axon-instruments-patch-clamp-amplifiers |
| Digidata 1440A digitizer | Molecular Devices | 1-2500-0173 D |
| Inline Solution Heater | Warner Instruments | SH-27B |
| SliceScope Pro 2000 microscope | Scientifica | https://www.scientifica.uk.com/products/slicescope-pro-2000 |

### RESOURCE AVAILABILITY

#### Lead contact

Further information and requests for resources and regents should be directed to and will be fulfilled by the lead contact, Daniel J Liebl

#### Materials available

This study did not generate new reagents.

#### Data and code availability

- Raw data is deposited in the Open Data Commons for Traumatic Brain Injury (ODC-TBI) database. RNA sequencing data is available in the Gene Expression Omnibus (GEO) repository for mouse scRNA-seq. and de-identified human bulk-seq.
- All code was implemented using existing Seurat pipelines.
- Any additional information needed to reanalyze the data reported is available from lead contact upon request.

### EXPERIMENTAL MODEL AND STUDY DETAILS

#### Sex as a Biological Variable

Our studies examined male and female mice, and sex-dependent effects are reported.

#### Animal

Mice between 2-4 months of age were used for all experiments. Wildtype mice (Jackson Laboratory, Bar Harbor, ME, #000664) and experimental mice were maintained on a C57BL/6J background. *Tmem119^CreERT2^:SRR^fl/fl^*and *CamKII^CreERT2^:Grin2b^fl/fl^* mice were generated by crossing *SRR^fl/fl^*^75^ and *Grin2b^fl/fl^ (*C57BL/6J-*Grin2b^em5Lutzy^*/J; Jackson Laboratory, #032664) mice to cell-specific cre expressing microglial *Tmem119^CreERT2^* (C57BL/6-*Tmem119 em2(EGFP)Gfng* /J; Jackson Laboratory, #031820) and neuronal *CamKII^CreERT2^ (*B6;129S6-Tg(Camk2a-cre/ERT2)1Aibs/J; Jackson Laboratory, #012362) mice, respectively. All animals were group-housed with a 12/12-hour light/dark cycle and had access to food and water *ad libitum*.

Cre recombinase expression was induced by six intraperitoneal (i.p.) injections of 90 mg/kg tamoxifen (Sigma Aldrich, St. Louis, MO, #T5648) beginning at least 2 weeks prior to experimentation for 1 day, followed by 2 days of rest, followed by 5 consecutive days of injection. Tamoxifen is prepared by dissolving 1 g in 20 mL of 90% sunflower seed oil (Sigma-Aldrich #S5007) and 10% ethanol. All mice excluding wildtype mice received tamoxifen treatment; *CamKII^CreERT2^:Grin2b^fl/fl^* mice began injections exactly 19 days prior to sham or CCI injury to minimize baseline differences in learning and memory due to prolonged NMDAR deficiencies.

#### Study Approval

All studies involving animal use were approved by the University of Miami Animal Use and Care Committee (#23-035), and the collection of human TBI tissues was approved by the University of Miami Institutional Review Board (#20080609).

#### Controlled Cortical Impact (CCI) injury model

CCI injury was performed as previously described^5–7,76,77^. Briefly, mice were anesthetized with a cocktail of ketamine/xylazine (100/10 mg/kg) in PBS, placed in a stereotactic frame, and a 5 mm craniotomy was made over the right parieto-temporal cortex (−2.0 mm A/P, 2.5 mm lateral from Bregma). A 3 mm pneumatic piston was used to produce a CCI injury (velocity: 4 m/s, depth:

0.55 mm, duration: 150 ms with an eCCI-6.0 device (Custom Design & Fabrication, Brookings, SD).

#### PSVue administration

A stock solution of 1 mM PSVue-480 (PSVue; Molecular Targeting Technologies, West Chester, PA, #P-1003) was prepared according to the manufacturer’s instructions. PSVue was further diluted in DMSO to a working concentration of 0.168 mg/mL and administered for 3 dpi using Alzet osmotic pumps (Alzet, #0000289).

#### Pharmacological agents

Ro 25-6981 maleate (Ro; Bio-Techne Corporation, Minneapolis, MN, #15-941-R) was administered for 3 dpi using Alzet osmotic pumps^7^ (Alzet, Cupertino, CA, #0000290). Ro was dissolved in 0.9% saline to final working concentrations of 1 nM, 10 nM, and 100 nM and preloaded into Alzet pumps one day before surgery; 0.9% saline was used as a vehicle control. Preloaded pumps were connected to a brain infusion cannula (Alzet, #0008851) and placed in 0.9% saline at 37°C to equilibrate overnight. Immediately following sham or CCI surgery, a stereotactic holder was used to place the infusion device contralateral to the injury site, so that the tip of the cannula reached the lateral ventricle (0.5 mm A/P, 1.5 mm lateral, −3.0 mm ventral from Bregma). The cannula was attached to the skull using Loctite 454 Prism Gel (#233998, Henkel Corp., Bridgewater, NJ) and the pump placed subcutaneously in the back of the neck.

### BEHAVIORAL METHODS DETAILS

#### Fear conditioning analysis

Mice were tested for contextual fear behavior at 5-7 days post injury (dpi) as previously described^7^. The test apparatus consisted of an isolation cubicle (Coulbourn Instruments, #H10-24T) with a fear conditioning chamber (Coulbourn Instruments, Allentown, PA, #H11-01M) connected to an electric grid floor. Mice were exposed to a 0.5 mA foot shock (Coulbourn Instruments, #H13-15) at 6 dpi, and percent freezing at 7 dpi was quantified using the video-based analysis software FreezeFrame 5 (Actimetrics).

#### Light-dark transition analysis

Mice were tested for anxiety-based behavior at 6 dpi. The testing chamber consisted of an arena (dimensions, in centimeters (cm): 37.5 (width), 26 (depth), 20 (height)) with two connected chambers, where one is brightly lit with white walls and the other is dimly lit with black walls. At the beginning of the test, the entryway between both chambers was blocked with a removable door, and mice were placed in the dark chamber for 1 minute. Then, the door was removed, and mice were allowed to explore freely between the two chambers for 10 minutes. Movement was recorded and analyzed for time spent in each chamber during the last 10 minutes of the test using the EthoVisionXT software (Noldus, Wageningen, the Netherlands).

#### Open field analysis

At 7 dpi, mice were placed in an open field arena (dimensions, in cm: 72 (width), 72 (depth), 40 (height) and allowed to freely explore for four 5-minute trials. The arena was cleaned with 70% ethanol between trials, and mice were returned to their home cage for a 20-minute break between trials. Movement was recorded and analyzed for total distance travelled and average velocity using the EthoVisionXT software (Noldus).

### METHODS DETAILS

#### Electrophysiology

At 1-week post-injury animals were decapitated under anesthesia with 4% isoflurane, 70% N_2_O, and 30% O_2_. The brain was quickly removed and chilled in 4⁰C sucrose-based artificial cerebrospinal fluid (sucrose-aCSF, in mM: 110 Sucrose, 5 D-glucose, 28 NaHCO_3_, 60 NaCl, 3 KCl, 1.25 NaH_2_PO_4_, 7 MgCl_2_, 0.5 CaCl_2_,) oxygenated with 95% O_2_ and 5% CO_2_. Brains were hemisected, the ipsilateral hippocampus was carefully dissected, and sectioned at 400μm thickness in oxygenated sucrose-based aCSF at 4⁰C using a Leica VT 1200 Vibrating Blade Microtome. Sectioned slices were transferred to oxygenated 50:50 sucrose-based aCSF and standard aCSF (in mM: 10 D-glucose, 25 NaHCO_3_, 125 NaCl, 2.5 KCl, 1.25 NaH_2_PO_4_, 1 MgCl_2_, 2 CaCl_2_) at room temperature for 20 minutes, then transferred to standard aCSF for 1 hour incubation at room temperature ^78,79^.

Slices were placed in a submerged recording chamber continuously perfused with oxygenated standard aCSF and heated to 31⁰C using an inline solution heater (Warner Instruments) and allowed to incubate for 20 minutes. A Platinum-iridium recording electrode was inserted to stimulate Schaffer collaterals and a recording pipette filled with 2M NaCl solution 1-2MΩ input resistance was placed in the stratum radiatum CA1 axonal projections to record field Excitatory Post-Synaptic Potentials (fEPSPs). Recorded signal was amplified using a Multiclamp 700B amplifier (Molecular Devices). The amplified signal was digitized at 20 kHz sampling frequency and low-pass filtered at 2 kHz using a Digidata 1440A and pClamp software (Molecular Devices)^80^.

Input-Output curve responses were obtained using a 20 μA step current until maximum fEPSP slope but not exceeding 200 μA. The stimulus current which elicited 40% of the maximum fEPSP slope was selected as the current for baseline stimulation at 0.033 Hz, paired-pulse facilitation response, high-frequency stimulation (HFS), and post-HFS response. A 20-minute baseline was obtained during which slices showing more than 10% variation in their normalized fEPSP slope were discarded due to instability. Paired pulse facilitation was tested at 10, 25, 50, 100, 200, and 400 ms pulse intervals. Long-term Potentiation (LTP) was induced via HFS consisting of a single train of 100 Hz over 1 second. Post-HFS response was recorded at 0.033 Hz for 60 minutes after HFS ^80^.

#### Multiplex panel

Mice were anesthetized at 3 dpi using a ketamine/xylazine cocktail and the hippocampus ipsilateral to the injury site harvested immediately. Tissues were frozen on dry ice and processed using the MILLIPLEX Mouse Cytokine/Chemokine Magnetic Bead Panel (Sigma-Aldrich #MCYTOMAG-70K-PMX) and Mouse MMP Magnetic Bead Panel 3 (Sigma-Aldrich #MMMP3MAG-79K).

#### Golgi staining

Mice were anesthetized at 3 or 7 dpi and whole brains harvested immediately. Brains were rinsed in distilled water and processed with the FD Rapid GolgiStain Kit according to the manufacturer’s protocol (FD NeuroTecchnologies, Columbia, MD, #PK401). For Ro-treated WT and female *SRR^fl/fl^* and *Tmem119^CreERT2^:SRR^fl/fl^* mice, 0.5 mm z-stack sections were imaged on a Zeiss Axiovert 200M fluorescence microscope with a 100X oil-immersion objective and 1.25X zoom. For *Grin2b^fl/fl^*, *CamKII^CreERT2^:Grin2b^fl/fl^,* and non-Ro treated WT mice, 0.2 mm z-stack sections were imaged on a Keyence fluorescence microscope (Keyence, Osaka, Japan, #BZ-X810) with a 100X oil-immersion objective and 1.2X zoom. For all animals, three dendritic segments (>10μm in length) in the CA1 region of the hippocampus were imaged per cell, where three cells were imaged per animal for a total of nine segments per animal. The Reconstruct software (Harris Lab, Center for Learning and Memory, University of Texas, Austin, TX) was used for analysis as previously described^29^.

#### Immunohistochemistry

Mice were anesthetized with a ketamine/xylazine cocktail and transcardially perfused with 1X PBS followed by 4% paraformaldehyde (PFA; ThermoFisher, Waltham, MA, #A11313.36) in 1X PBS (pH 7.4). Brains were dissected and postfixed in 4% PFA for 24 hours at 4°C, then transferred to 30% sucrose (VWR #0355) in 0.01% sodium azide for at least 48 hours. Brains were embedded in a 50% OCT (VWR #4583) and 50% sucrose solution, frozen using isopropyl alcohol, and sectioned at 30 mm using a Leica CM1900 cryostat.

For PSVue immunohistochemistry, sections were warmed to room temperature (RT) and incubated in 1X PBS for 10 minutes to remove excess OCT, then washed 3 times for 15 minutes in 0.4% PBS-Triton-X followed by blocking in 5% normal goat serum (NGS) in 0.4% PBS-Triton-X for 1 hour at RT. Sections were incubated overnight at 4°C with the following primary antibodies (1:500 in blocking solution): rabbit anti-PSD-95 (ThermoFisher, #51-8900) and Guinea pig anti-Vglut1 (Synaptic Systems, Göttingen, Germany, #135304). Sections were warmed to RT and washed 3 times for 15 minutes in 0.1% PBS-Triton-X followed by a 2-hour incubation with secondary antibodies (1:500 in 0.1% PBS-Triton-X): Alexa Fluor^TM^ 647 goat anti-rabbit (ThermoFisher, #1524909) and Alexa Fluor^TM^ 594 goat anti-guinea pig (ThermoFisher, #2540867). Sections were washed 3 times for 15 minutes in 0.1% PBS-Triton-X and cover-slipped with Fluoro-Gel mounting media (Electron Microscopy Sciences, Hatfield, PA, #17985-10) with DAPI (1:1000). For C1q staining, an additional blocking step was performed to block endogenous IgG due to higher background levels. Following blocking in 5% NGS, sections were incubated with 40 mg/mL unconjugated AffiniPure Fab Fragment Goat αMouse IgG (H+L) (Jackson Immuno, West Grove, PA, #715-007-003) in 1X PBS for 2 hours at RT. The same protocol as above was followed for incubation with primary and secondary antibodies: rabbit anti-PSD-95, guinea pig anti-Vglut1, rat anti-C1qa (Novus Biologicals, Centennial, CO, #NB200-539), Alexa Fluor^TM^ 488 goat anti-rabbit (ThermoFisher, #A11008), Alexa Fluor^TM^ 594 goat anti-guinea pig, and Alexa Fluor^TM^ 647 goat anti-rat (ThermoFisher, #A21247). After washing in 0.1% PBS-Triton-X, sections were incubated for 5 minutes at RT with 0.15% Sudan Black B (Sigma Aldrich, #199664) prepared in 70% ethanol^81^. Sections were washed in 70% ethanol for 30 seconds then 1X PBS for 5 minutes before cover-slipping.

For all tissues, three sections ∼100 mM apart per animal in the *SR* and *SLM* in the CA1 region of the hippocampus were imaged on a Dragonfly high speed confocal microscope (Oxford Instruments, Abingdon, England) using a 63X oil immersion objective and 0.2 μM step size.

#### Imaris Analysis

Images were deconvoluted using ClearView-GPU Deconvolution (Oxford Instruments) and analyzed with Imaris 10.0 software (Oxford Instruments). All visible Vglut1 and PSD95 puncta were identified using the Imaris “spots” function and quality filtering, and Vglut1 positive spots that colocalized with, or were within 0.5 μM of, PSD95 positive spots were defined as synapses. Areas of the *SR* and *SLM* were manually contoured using the “surfaces” function, and synapses were filtered for spots that were localized within these regions of interest. Synapses were then filtered for colocalization with PS or C1q, and the number of PS or C1q positive synapses was divided by the total number of synapses within the area of interest to obtain the percent of PS or C1q positive synapses. Average values from three sections per animal per region of interest were obtained.

#### Western blots analysis

Mice were anesthetized and perfused with 1X PBS. For whole tissue lysates, hippocampi ipsilateral to the injury site were dissected and lysed in 500 μL RIPA buffer with 1X protease (Sigma Aldrich, #04693159001) and phosphatase inhibitors (Sigma Aldrich, #04906845001). Samples were incubated on ice for 20 minutes, centrifuged at 4°C for 10 minutes (13.2g), and the pellet discarded. For isolated plasma membrane samples, hippocampi were processed with the Plasma Membrane Protein Isolation and Cell Fractionation kit (Invent Biotechnologies, #SM-005) according to the manufacturer’s instructions and finally resuspended in RIPA buffer with 1X protease and phosphatase inhibitors. For human samples, protein was obtained during RNA extraction using the Direct-zol RNA Miniprep kit (Zymo Research, Irvine, CA, #R2051) according to the manufacturer’s instructions and resuspended in RIPA buffer with 1X protease and phosphatase inhibitors. Control cortical protein lysates were obtained from cardiac arrest cases (Novus Biologicals, #NB820-59186, #NB820-60574, #NB820-59193, #NB820-59198, and #NB820-59177).

All samples were further processed with the Quick Start Bovine Serum Albumin (BSA) Standard kit (Bio-Rad, Hercules, CA, #5000206) to determine protein concentration. Samples were diluted to 30 μg and mixed with loading buffer composed of 90% NuPAGE^TM^ LDS Sample Buffer (Thermo Fisher, #NP0007) and 10% 2-Mercaptoethanol (Sigma Aldrich, M6250). Samples were separated on a 4%–20% stain-free gradient gel (Bio-Rad, Hercules, CA, #5678094 and #5678095), imaged on a ChemiDoc MP Imaging System (BioRad, version 2.3.0.07), and transferred to a nitrocellulose membrane using the Trans-Blot Turbo RTA Transfer Kit (Bio-Rad, #1704270). Total protein transferred was imaged and membranes were blocked for 1 hour at RT with 5% bovine serum albumin (BSA) in 1X PBS followed by overnight incubation at 4°C with primary antibodies diluted in 5% BSA (1:1000 unless otherwise noted): rabbit anti-BDNF (Abcam, Boston, MA, #ab9793), rabbit anti-phospho-ERK1/2 (1:500; Cell Signaling, Danvers, MA, #9101S), rabbit anti-ERK1/2 (Cell Signaling #9102), rabbit anti-NR1 (Millipore #05-432), rabbit anti-NMDAR1 (Abcam #ab109182), rabbit anti-NR2A (Millipore #07-632), rabbit anti-NR2B (Millipore #06-600), rabbit anti-GRIN2B (1:500; ThermoFisher, #21920-1-AP), rabbit anti-DAPK1 (Cell Signaling, Danvers, MA, #3008S), Rabbit anti-p-CREB (ABclonal, Woburn, MA, #AP0019), rabbit anti-CREB (ABclonal, #A11064), rabbit anti-Sodium Potassium ATPase (1:500; Abcam, #AB76020), rabbit anti-Phospho-N2B,Ser1303 (Millipore, #07-398), rabbit anti-C1qA (Abcam, #ab189922), rabbit anti-C3 (Invitrogen, #PA5-21349), mouse anti-GPR56 (Biolegend, #358205), mouse anti-SRR (Santa Cruz Biotech, Dallas, TX, #sc-365217), rabbit anti-Slc1a4 (Cell Signaling, #8442S), and mouse anti-GAPDH (1:5000, Santa Cruz Biotech, #47724). Membranes were then washed 3 times 15 minutes at RT in 1X TBS (Bio-Rad, #1706435) with 0.1% Tween-20 and incubated for 2 hours with respective secondary antibodies diluted 1:5000 in 5% BSA: goat anti-rabbit (Jackson Immuno, #111-035-003) and goat anti-mouse (Jackson Immuno, #115-035-003). Membranes were exposed with SuperSignal West Pico PLUS Chemiluminescent Substrate (ThermoFisher, #34578) and imaged using a ChemiDoc Touch Imaging System (Version 1.2.0.12, Bio-Rad). Image Lab software (Bio-Rad) was used to perform densitometry analysis, where protein measurements were normalized to GAPDH for human samples and total protein for mouse samples.

#### Fluorescence-Activated Cell Sorting (FACS)

Mice were anesthetized, perfused with 1X PBS, and ipsilateral hippocampi were extracted and processed with the Adult Brain Dissociation Kit (Miltenyi Biotech, Bergish Gladbach, Germany, #130-107-677) following the manufacturer’s instructions. Briefly, tissues were placed in a solution of pre-warmed enzymes 1 and 2, dissociated with a gentleMACS™ Octo Dissociator (Miltenyi Biotech, #130-096-427), and strained over a 70 μm filter in cold Hibernate A solution without calcium (Thermo Fisher, #NC0176976). Myelin debris removal was performed and the cell pellet was labelled with primary antibodies diluted 1:50 in flow buffer for 1 hour at 4°C: CD45-PerCP-Vio 700, REAfinity™ (Miltenyi Biotech, #130-110-801), CD11b-APC (Biolegend, San Diego, CA, #101212), ACSA2-PdE (Miltenyi, #130-116-141), and GPR56-PE/Cy7 (Biolegend, #358205). DAPI (0.1mg/mL; Miltenyi #130-111-570) was added to each sample, and cells were sorted on a MoFlo Astrios Cell Sorter (Beckman Coulter, Brea, CA).

#### Flow Cytometry

A similar protocol was followed as described above, however prior to primary antibody labelling samples were incubated with FcR blocking reagent TruStain FCX™ PLUS (1:100; Biolegend, #156603) in flow buffer for 5 minutes at RT. Incubation with CD45, CD11b, and ACSA-2 antibodies was performed as described above, following which cells were washed in flow buffer and fixed with 1% PFA in permealization buffer (0.5% Tween-20 in 1X PBS) for 1 hour at RT. Cells were resuspended in permealization buffer, and incubated with a mouse anti-SRR antibody (1:500; BD Biosciences, Franklin Lakes, NJ, #612053) overnight at 4°C. Samples were then incubated with a goat anti-mouse 488 secondary antibody (1:500; Thermo Fisher, #A1101) for 30 minutes at RT, resuspended flow buffer, treated with NucBlue Fixed Cell DAPI (Thermo Fisher, #R37606), and analyzed with a Cytoflex S flow cytometer (Beckman Coulter) and CytExpert software (Beckman Coulter). Fluorescent signal from unstained and single channel control samples was used to determine gating parameters, where astrocytes express ACSA-2 and microglia express high levels of CD11b and low levels of CD45 relative to CD45^high^ macrophages.

#### Parse scRNA-seq

Samples were dissociated into single cell suspensions similar to the FACS procedure described above, with the following modifications. To prevent transcriptional activation during dissociation, actinomysin D (5 μg/mL; Sigma Aldrich, #A1410), triptolide (10μM; Sigma Aldrich, #T3652), and anisomycin (27.1 μg/mL; Sigma Aldrich, #A9789) were added to the dissociation buffers^82^. In addition, RNAsin Plus RNase Inhibitor (Promega, #N2615) was added to the dissociation buffers to preserve RNA integrity. Myelin debris removal was performed twice, cells were resuspended in 1 mL of 5% BSA Fraction V (ThermoFisher, #15260037) in 1X PBS, and assessed for viability using Acridine orange & Propidium iodide (AO/PI). Samples with over 90% live cells were processed further with the EvercodeTM Cell Fixation v2 kit (Parse Biosciences, Seattle, WA). To increase cell retention, 15 mL Falcon tubes used during fixation were coated with 1% BSA Fraction V per the manufacturer’s recommendations. Samples were frozen using a Mr. FrostyTM container (ThermoFisher, #5100-001) and stored in −80°C until processing with the EvercodeTM Mega v2 kit (Parse Biosciences) according to the manufacturer’s protocol.

Sequencing libraries were sequenced on the NovaSeqX Plus targeting 50,000 reads per cell. Library demultiplexing, alignment, and gene quantification was performed using default parameters with the Parse Biosciences Pipeline (split pipe version 1.1.2). Resulting gene expression matrixes were processed using Seurat v5 calling the ‘ReadParseBio’ function. Downstream quality control, normalization, and data integration using CCA was implemented in Seurat v5^83^. Cell types were determined based on known cell type markers, and differential gene expression analysis was performed using the MAST test implemented in Seurat v5 (https://genomebiology.biomedcentral.com/articles/10.1186/s13059-015-0844-5). Microglia, including dividing and non-dividing microglia groups, and astrocyte cell types were further subclustered using the standard Seurat v5 pipeline with a resolution of 0.2. Differentially expressed genes were filtered based on a Log_2_FC ≥ +0.3 or ≤ −0.3 and adjusted p-value <0.05 and processed for GO:CC and GO:BP pathway analysis using GProfiler^84^.

#### Bulk RNA sequencing

RNA extraction was performed using the Direct-zol RNA Miniprep kit (Zymo Research, #R2051) and samples were shipped on dry ice to Azenta Life Sciences (South Plainfield, NJ) for RNA-sequencing. Control cortical RNA samples were obtained from cardiac arrest cases (Takara Bio, Kusatsu, Japan, #636570, #636571, and #636564). Libraries were prepared with rRNA depletion and paired end (PE) reads were sequenced on an Illumina NovaSeq. FASTQ files were processed using TrimGalore for trimming adapters and STAR (v2.5.2) for alignment to the human genome (hg38). Gene quantification against GENCODE (v35) was performed using the geneCounts function in STAR. Batch correction for two groups that were sequenced on different dates was performed using ComBat-seq^85^, and differential expression was determined using the R package DESeq2. Differentially expressed genes were filtered based on a Log_2_FC ≥ +2 or ≤ −2 and adjusted p-value <0.05 and processed for GO:CC and GO:BP pathway analysis using GProfiler^84^

### STATISTICAL ANALYSIS

Data was analyzed using a one- or two-way ANOVA with Tukey’s multiple comparisons or a Student’s two-tailed *t-*test using GraphPadPrism 9 (San Diego, CA). In the case of a non-normal distribution, a Mann-Whitney test was used. All graphs represent mean values, with error bars depicting the standard error of the mean (SEM) and *, **, and *** representing *p*-values equal to or less than 0.05, 0.01, and 0.001 respectively.

