## Supplementary material for "Cognitive dysfunction following brain trauma results from sex-specific reactivation of the developmental pruning processes": All supplemental data

**SUPPLEMENTAL DATA**
**Table S1**

| Sub-cluster | Sex | GO: PW | Gene # | GO description |
|---|---|---|---|---|
| A0 (Sham vs CCI) | Male up | BP | 3 | Synapse pruning |
| | | | 3 | Complement activation, Classical pathway |
| | | | 3 | Cell junction disassembly |
| | | | 3 | Complement activation |
| | | CC | 3 | Complement component C1 complex |
| | | | 3 | Complement component C1q complex |
| | Male down | BP | 9 | Regulation of synapse organization |
| | | | 9 | Regulation of synapse structure or activity |
| | | | 11 | Synapse organization |
| | | | 13 | Cell junction organization |
| | | CC | 15 | Glutamatergic synapse |
| | | | 21 | Synapse |
| | | | 22 | Cell junction |
| | | | 13 | Post-synapse |
| | | | 12 | Dendrite |
| | | | 12 | Dendritic tree |
| | | | 17 | Neuron projection |
| | | | 16 | Plasma membrane region |
| | | | 14 | Somatodendritic compartment |
| | | | 21 | Cell projection |
| | Female up | BP | 6 | Cell junction |
| | | CC | 5 | Post-synaptic specialization |
| | | | 4 | Postsynaptic membrane |
| | | | 3 | Presynaptic active zone |
| | | | 3 | GABAergic synapse |
| | | | 7 | Neuron project |
| A1 (Sham vs CCI) | Male up | BP | 11 | Cell junction |
| | | | 3 | Synapse pruning |
| | | | 3 | Complement activation, classical pathway |
| | | | 3 | Cell junction disassembly |
| | | CC | 3 | Complement component C1 complex |
| | | | 3 | Complement component C1q complex |
| A2 (CCI vs CCI) | Female up | CC | 27 | Cell projection |
| | | | 26 | Plasma membrane bounded cell projection |
| | | | 19 | Neuron projection |
| | | | 16 | Somatodendritic compartment |
| | | | 13 | Axon |
| | Female up | BP | 20 | Neurogenesis |

| | | | | |
|---|---|---|---|---|
| A3 (Sham vs CCI) | | CC | 13 | Synapse |
| | | | 8 | Glutamatergic synapse |
| | | | 14 | Neuron projection |
| | | | 16 | Plasma membrane bounded cell proj. |
| | | | 9 | Neuronal cell body |
| | | | 9 | Cell body |
| | | | 10 | Somatodendritic compartment |
| | | | 14 | Cell junction |
| | | | 2 | Calcium- and calmodulin-dependent protein kinase complex |
| | | | 17 | Cell projection |
| | Female down | BP | 14 | Modulation of chemical synaptic transmission |
| | | | 14 | Regulation of trans-synaptic signaling |
| | | | 14 | Chemical synaptic transmission |
| | | | 14 | Anterograde trans-synaptic signaling |
| | | | 14 | Trans-synaptic signaling |
| | | | 14 | Synaptic signaling |
| | | CC | 15 | Synapse |
| | | | 7 | Postsynaptic membrane |
| | | | 8 | Synaptic membrane |
| | | | 19 | Cell junction |
| | | | 17 | Neuron projection |
| | | | 10 | Postsynapse |
| | | | 19 | Cell projection |
| | | | 15 | Synapse |
| A4 (Sham vs CCI) | Male up | BP | 7 | Cellular component organization |
| | | CC | 2 | Postsynaptic actin cytoskeleton |
| | | | 1 | Postsynaptic cytoskeleton |
| | Male down | CC | 2 | Synapse-associated extracellular matrix |
| A5 (Sham vs CCI) | Male down | BP | 6 | Synapse organization |
| | | | 5 | Regulation of synapse organization |
| | | | 5 | Regulation of synapse structure or activity |
| | | CC | 7 | Synaptic membrane |
| | | | 9 | Synapse |
| | | | 8 | Plasma membrane region |
| | | | 5 | Post-synaptic membrane |
| | | | 6 | Glutamatergic synapse |
| | | | 9 | Cell junction |
| | | | 6 | Postsynapse |
| | | | 2 | Synapse-associated extracellular matrix |
| | | | 2 | Postsynaptic membrane |
| | | | 3 | GABA-ergic synapse |
| | | | 4 | Postsynaptic density |
| | | | 2 | Extrinsic component of synaptic membrane |
| | | | 4 | Asymmetric synapse |

| | | | | |
|---|---|---|---|---|
| | | | 4 | Postsynaptic specialization |
| | | | 4 | Neuron to neuron synapse |

| | | | | |
|---|---|---|---|---|
| M0 (Sham vs CCI) | Male up | BP | 21 | Cell junction organization |
| | | | 12 | Translation to synapse |
| | | | 12 | Translation at postsynapse |
| | | | 12 | Translation at synapse |
| | | CC | 48 | Synapse |
| | | | 35 | Post-synapse |
| | | | 23 | Presynapse |
| | | | 16 | Neuron to neuron synapse |
| | | | 15 | Asymmetric synapse |
| | | | 14 | Postsynaptic density |
| | | | 14 | Postsynaptic specialization |
| | | | 17 | Glutamatergic synapse |
| | | | 10 | Schaffer collateral - CA1 synapse |
| | | | 19 | Neuronal cell body |
| | | | 21 | Cell body |
| | | | 26 | Somatodendritic compartment |
| | | | 61 | Cell junction |
| | Female down | BP | 15 | Phagocytosis |
| | | | 5 | Synaptic pruning |
| | | | 5 | Cell junction disassembly |
| | | CC | 48 | Neurogenesis |
| | | | 39 | Synapse |
| | | | 3 | Complement component C1 complex |
| | | | 3 | Complement component C1q complex |
| | | | 56 | Cell junction |
| | | | 12 | Cell-substrate junction |
| | | | 4 | Glycinergic synapse |
| M2 (Sham vs CCI) | Male up | CC | 27 | Synapse |
| | | | 31 | Cell junction |
| | | | 19 | Somatodendritic compartment |
| M3 (Sham vs CCI) | Male up | BP | 12 | Translation at postsynapse |
| | | | 12 | Translation at synapse |
| | | | 11 | Translation at presynapse |
| | | | 24 | Regulation of cell projection organization |
| | | | 14 | Regulation of synapse organization |
| | | | 20 | Synapse organization |
| | | | 25 | Cell junction organization |
| | | CC | 77 | Synapse |
| | | | 24 | Glutamatergic synapse |
| | | | 27 | Presynapse |
| | | | 43 | Postsynapse |

| | | | | |
|---|---|---|---|---|
| | | | 63 | Cell projection |
| | | | 41 | Neuronal projection |
| | | | 25 | Dendrite Tree |
| | | | 12 | Dendritic spine |
| | | | 12 | Neuron spine |
| | | | 19 | Cell-cell junction |
| | | | 10 | Cell substrate junction |
| | | | 16 | Neuron-neuron synapse |
| | | | 16 | Asymmetric synapse |
| | | | 23 | Anchoring junction |
| | | | 25 | Dendrite |
| | | | 40 | Somatodendritic compartment |
| | | | 90 | Cell junction |

**GO synaptic-related pathways observed in male and female astrocyte and microglia populations.** Comparison between sham and controlled cortical impact (CCI) conditions were compared for all subclusters showing a significant upregulation and downregulation of synaptic-related pathways. Subcluster labels, sex, directional change, GO pathway (GO:PW) for cellular component (CC) and biological processes (BP), number of genes changed, and the original GO pathway are all shown. Gene list and pathway consolidation list can be found on the associated with supplemental file 1.

**Table S2.**

| Astrocytes | | | | Microglia | |
|---|---|---|---|---|---|
| **Male up** | **Male down** | **Female up** | **Female down** | **Male up** | **Female down** |
| ACTB | ADAM23 | ACTB | ABR | ABR | ABCC1 |
| AHNAK | ANKS1 | ADGRL3 | ALDH1A1 | ABCG1 | ARF1 |
| APOBEC1 | APLP2 | AGBL4 | ATP2B2 | ANXA2 | ARHGDIA |
| C1QA | ATP1A2 | BMPR1B | BCAR3 | AP1S2 | ARRB1 |
| C1QB | BCAN | BTBD3 | BRINP3 | AP3S1 | ATP6AP1 |
| C1QC | CHD4 | CAMK2A | CNTN1 | AFAP1 | B2M |
| CAMK1D | CPEB4 | CDH6 | DCC | AKR1A1 | B4GALT1 |
| CSF1R | DAG1 | CPE | DOCK4 | APOB | BTG2 |
| FAM107A | DGKB | CTNNA2 | EPHA4 | APOD | C1QA |
| GJB6 | EPS15 | DNER | EPHB1 | ARHGEF7 | C1QB |
| LGMN | EPS8 | ENPP2 | FAM107A | ARRB2 | C1QC |
| MKI67 | FAT3 | FABP7 | KCNQ3 | ATP6V0A2 | C5AR1 |
| MYO5A | FERMT2 | GABRA2 | KIFC3 | BAIAP2 | CALM2 |
| NFASC | GABRG1 | GPM6B | GJB6 | CD63 | CALR |
| NRXN3 | GPM6A | GRIN3A | GLI2 | CD86 | CANX |
| SPARC | GRHL1 | GRIP1 | GLUL | CLDN11 | CBL |
| TMSB4X | GRIA1 | HES5 | GM14569 | CTSL | CD47 |
| VCL | GRK3 | HTR2C | GPHN | DBNL | CD53 |
| VIM | KCNJ10 | KCND2 | GPR158 | DGKI | CDC42 |
| | IFT57 | LEF1 | GRIN2C | DNAJC13 | CDH23 |
| | LGI1 | LGI4 | GSTM5 | EPB41L3 | CDK5RAP2 |
| | LYPD6 | MMD2 | PLCB1 | FLCN | CERS2 |
| | MAGI2 | MYO6 | PLCL1 | FMN1 | CLCN4 |
| | MFN1 | NR2F2 | PREX1 | FYN | CSF1R |
| | NEGR1 | PBX1 | PSD2 | GABBR1 | CYSLTR1 |
| | OGT | RGS9 | PTPRD | GDI2 | CYBA |
| | PTPRD | SEMA6D | RGS6 | GLUL | DDX6 |
| | RPS6KB1 | SLC7A11 | SLC7A10 | GNAS | EIF3D |
| | SLC13A5 | SNTB1 | SLC24A4 | GRN | ENTPD1 |
| | SLC6A11 | SPARCL1 | | H2-D1 | EP300 |
| | SLCO1C1 | TENM4 | | HNRNPA2B1 | FCER1G |
| | SPARCL1 | TNC | | HIPK1 | GLUL |
| | SYNE1 | TRIM9 | | IL10RA | GPHN |
| | VCAM1 | UNC5C | | IL15RA | HDAC4 |
| | | | | IFI30 | HEXB |
| | | | | KNL1 | HNRNPK |
| | | | | LAMP1 | HS6ST1 |
| | | | | MPP1 | HSPA5 |
| | | | | MYO5A | HSPA8 |
| | | | | NAIP6 | IFNGR1 |
| | | | | NCF1 | IL16 |
| | | | | NECTIN2 | ITGAV |
| | | | | NFASC | ITGB1 |
| | | | | PABPC1 | ITGB3 |
| | | | | PICALM | ITM2C |

| | | | | | |
|---|---|---|---|---|---|
| | | | | PIK3C3 | ITPR1 |
| | | | | PLCB2 | JADE2 |
| | | | | PLEKHG5 | JAM3 |
| | | | | PPFIA4 | KLF7 |
| | | | | PSAP | LIFR |
| | | | | RAB11A | MED12 |
| | | | | RAB32 | MSN |
| | | | | RACK1 | MYO7A |
| | | | | RFTN1 | PLXNB2 |
| | | | | RPL7A | PPT1 |
| | | | | RPL9 | PRAG1 |
| | | | | RPL11 | PRKCSH |
| | | | | RPL12 | PTPN6 |
| | | | | RPL13A | PURA |
| | | | | RPL17 | PXN |
| | | | | RPL21 | RAC1 |
| | | | | RPL28 | RGS2 |
| | | | | RPL32 | RHOA |
| | | | | RPL36A | RHOB |
| | | | | RPL37 | SERPINF1 |
| | | | | RPL38 | SIGLECE |
| | | | | RPS2 | SLC16A7 |
| | | | | RPS3 | SLC29A1 |
| | | | | RPS3A1 | SLC3A2 |
| | | | | RPS4X | SOX4 |
| | | | | RPS5 | SMURF1 |
| | | | | RPS7 | SPACDR |
| | | | | RPS11 | SPARC |
| | | | | RPS19 | SRGN |
| | | | | RPS21 | STON2 |
| | | | | RPS25 | STXBP1 |
| | | | | RPSA | SYNGR2 |
| | | | | SDCBP | SYPL |
| | | | | SH3BP1 | TLN1 |
| | | | | SLC27A1 | TGM2 |
| | | | | STAT3 | TLR4 |
| | | | | STK38 | TMED9 |
| | | | | STXBP2 | TMEM106B |
| | | | | SV2A | TMEM30A |
| | | | | TBC1D16 | TNFSF13B |
| | | | | TNR | TREM2 |
| | | | | TRPV4 | TYROBP |
| | | | | TUBB4A | UBB |
| | | | | VPS4B | WAS |
| | | | | ZNRF2 | XRCC5 |
| | | | | | YTHDF2 |

**Unique upregulated or downregulated genes associated with astrocytes or microglia from male and female mice.** Consolidated gene list from all astrocyte or microglia subclusters show uniquely expressed upregulated or downregulated genes associated with male or female mice.

**Table S3.**

| Sub-cluster | Sex | GO:PW | Gene # | GO description |
|---|---|---|---|---|
| A0 (Sham vs CCI) | $SRR^{fl/fl}$ up | BP | 4 | Synapse organization |
| | | | 4 | Cell junction organization |
| | | | 1 | Synapse pruning |
| | | | 1 | Complement activation, classical pathway |
| | | CC | 5 | Synapse |
| | | | 5 | Cell junction |
| | | | 1 | Complement component C1 complex |
| | | | 1 | Complement component C1q complex |
| | $SRR^{fl/fl}$ down | BP | 10 | Neurogenesis |
| | | CC | 10 | Dendrite |
| | | | 13 | Dendritic tree |
| | | | 16 | Neuron projection |
| | | | 16 | Plasma membrane bounded cell projection |
| | | | 15 | Cell projection |
| | TMEM119: $SRR^{fl/fl}$ down | BP | 3 | Regulation of neurotransmitter receptor localization to postsynaptic specialization membrane |
| | | | 3 | Regulation of protein localization to synapse |
| | | | 3 | Regulation of receptor localization to synapse |
| | | | 4 | Regulation of postsynaptic membrane neurotransmitter receptor levels |
| | | | 3 | Protein localization to postsynaptic specialization membrane |
| | | | 3 | Neurotransmitter receptor localization to postsynaptic specialization membrane |
| | | CC | 6 | Glutamatergic synapse |
| | | | 5 | Postsynaptic density |
| | | | 5 | Asymmetric synapse |
| | | | 5 | Postsynaptic specialization |
| | | | 8 | Synapse |
| | | | 5 | Neuron to neuron synapse |
| | | | 9 | Cell junction |
| | | | 6 | Postsynapse |
| A1 (Sham vs CCI) | $SRR^{fl/fl}$ up | BP | 8 | Synapse organization |
| | | | 9 | Cell junction organization |
| | | CC | 10 | Synapse |
| | | | 11 | Cell junction |
| | | | 7 | Postsynapse |
| | | | 1 | Complement component C1q complex |
| | | | 1 | Complement component C1 complex |
| | $SRR^{fl/fl}$ down | BP | 41 | Neurogenesis |
| | | | 30 | Cell projection organization |
| | | | 10 | Regulation of axonogenesis |

| | | | | |
|---|---|---|---|---|
| | | | 15 | Axonogenesis |
| | | | 18 | Regulation of cell projection organization |
| | | | 15 | Axon development |
| | | | 18 | Cell junction organization |
| | | CC | 35 | Cell junction |
| | | | 26 | Neuron projection |
| | | | 34 | Plasma membrane bounded cell projection |
| | | | 34 | Cell projection |
| | | | 26 | Synapse |
| | | | 17 | Postsynapse |
| | | | 12 | Synaptic membrane |
| | | | 7 | Adherens junction |
| | | | 9 | Postsynaptic membrane |
| A3 (Sham vs CCI) | SRR$^{fl/fl}$ up | CC | 7 | Cell projection |
| | | | 2 | Postsynaptic cytoskeleton |
| | | | 2 | Presynaptic cytosol |
| | | | 6 | Plasma membrane bounded cell projection |
| | | | 5 | Neuron projection |
| | | | 4 | Axon |
| | | | 2 | Cytosolic region |
| M0 (Sham vs CCI) | SRR$^{fl/fl}$ up | BP | 9 | Translation at presynapse |
| | | | 9 | Translation at postsynapse |
| | | | 9 | Translation at synapse |
| | | CC | 23 | Synapse |
| | | | 23 | Cell junction |
| | | | 18 | Postsynapse |
| | | | 11 | Presynapse |
| | | | 2 | Complement component C1q complex |
| | | | 2 | Complement component C1 complex |
| | SRR$^{fl/fl}$ down | BP | 33 | Neurogenesis |
| | | | 25 | Neuron projection development |
| | | | 19 | Neuron projection morphogenesis |
| | | | 9 | Regulation of axon extension |
| | | | 11 | Neuron projection extension |
| | | | 9 | Axon extension |
| | | | 13 | Axonogenesis |
| | | CC | 30 | Cell junction |
| | | | 25 | Neuron projection |
| | | | 21 | Somatodendritic compartment |
| | | | 18 | Axon |
| | | | 18 | Dendrite |
| | | | 18 | Dendritic tree |
| | | | 8 | Dendritic spine |
| | | | 8 | Neuron spine |

| | | | | |
|---|---|---|---|---|
| **M1 (CCI vs CCI)** | **SRR$^{fl/fl}$ up** | BP | 21 | Translation at postsynapse |
| | | | 21 | Translation at synapse |
| | | | 20 | Translation at presynapse |
| | | | 4 | Synapse pruning |
| | | | 4 | Complement activation, classical pathway |
| | | | 4 | Cell junction disassembly |
| | | CC | 50 | Synapse |
| | | | 37 | Postsynapse |
| | | | 51 | Cell junction |
| | | | 26 | Presynapse |
| | | | 13 | Postsynaptic density |
| | | | 13 | Asymmetric synapse |
| | | | 3 | Complement component C1 complex |
| | | | 3 | Complement component C1q complex |
| | | | 13 | Postsynaptic specialization |
| | | | 13 | Neuron to neuron synapse |
| | | | 17 | Somatodendritic compartment |
| **M2 (CCI vs CCI)** | **SRR$^{fl/fl}$ up** | BP | 46 | Translation at postsynapse |
| | | | 46 | Translation at synapse |
| | | | 45 | Translation at presynapse |
| | | CC | 166 | Synapse |
| | | | 179 | Cell junction |
| | | | 101 | Postsynapse |
| | | | 113 | Cell projection |
| | | | 84 | Presynapse |
| | | | 75 | Neuron projection |
| | | | 66 | Somatodendritic compartment |
| | | | 35 | Asymmetric synapse |
| | | | 16 | Phagocytic vesicle |
| | | | 36 | Neuron to neuron synapse |
| | | | 25 | Synaptic vesicle |
| | | | 32 | Postsynaptic density |
| | | | 34 | Postsynaptic specialization |
| | | | 9 | Autophagosome membrane |
| | | | 47 | Dendrite |
| | | | 47 | Dendritic tree |
| | | | 6 | Phagocytic cup |
| | | | 12 | Autophagosome |
| | | | 3 | Complement component C1q complex |
| | | | 3 | Complement component C1 complex |
| | | | 6 | Phagocytic vesicle membrane |
| | | | 15 | Synaptic vesicle membrane |
| **M3 (CCI vs CCI)** | **SRR$^{fl/fl}$ up** | BP | 39 | Translation at synapse |
| | | | 39 | Translation at postsynapse |
| | | | 38 | Translation at presynapse |
| | | | 5 | Synapse pruning |
| | | | 7 | Modification of synaptic structure |

| | | | | |
|---|---|---|---|---|
| | | | 6 | Modification of postsynaptic structure |
| | | CC | 121 | Cell junction |
| | | | 129 | Synapse |
| | | | 79 | Postsynapse |
| | | | 61 | Presynapse |
| | | | 25 | Asymmetric synapse |
| | | | 26 | Neuron to neuron synapse |
| | | | 25 | Postsynaptic specialization |
| | | | 23 | Postsynaptic density |
| | | | 31 | Glutamatergic synapse |
| | | | 43 | Somatodendritic compartment |
| | | | 32 | Neuronal cell body |
| | | | 3 | Complement component C1q complex |
| | | | 3 | Complement component C1 complex |
| | | | 71 | Cell projection |
| | | | 49 | Neuron projection |
| | | | 30 | Dendrite |
| | | | 30 | Dendritic tree |
| | | | 20 | Cell leading edge |
| **M6 (CCI vs CCI)** | **SRR$^{fl/fl}$ up** | BP | 3 | Synapse pruning |
| | | | 3 | Complement activation, classical pathway |
| | | | 3 | Cell junction disassembly |
| | | | 3 | Complement activation |
| | | CC | 3 | Complement component C1 complex |
| | | | 3 | Complement component C1q complex |
| | | | 7 | Synapse |
| | | | 5 | Postsynapse |
| | | | 7 | Cell junction |

**GO synaptic-related pathways observed in *SRR$^{fl/fl}$* and *Tmem119$^{creErt2}$:SRR$^{fl/fl}$* astrocyte and microglia populations.** Comparison between sham and controlled cortical impact (CCI) conditions were compared for all subclusters showing a significant upregulation and downregulation of synaptic-related pathways. Subcluster labels, sex, directional change, GO pathway (GO:PW) for cellular component (CC) and biological processes (BP), number of genes changed, and the original GO pathway are all shown. Gene list and pathway consolidation list can be found on the associated with supplemental file 1.

**Table S4.**

| Astrocytes | | | | Microglia | |
|---|---|---|---|---|---|
| **SRR$^{fl/fl}$ up** | **SRR$^{fl/fl}$ down** | **TMEM:SRR$^{fl/fl}$ up** | **TMEM:SRR$^{fl/fl}$ down** | **SRR$^{fl/fl}$ up** | **SRR$^{fl/fl}$ down** |
| ACTB | ABR | | CPEB4 | C1QB | ABI3 |
| ASIC2 | ACSL3 | | DGKB | C1QC | ABL1 |
| C1QA | ACSL6 | | EPS15 | CFL1 | ADGRG1 |
| C1QB | ADGRL1 | | GPC6 | EEA1 | ALOX5 |
| C1QC | AFAP1 | | GRID2 | EIF5B | AKNA |
| CDH19 | ANKS1 | | KALRN | MYO9A | ARHGAP22 |
| FABP7 | APBA2 | | LAMA3 | PABPC1 | ARRB1 |
| KCND2 | ARHGAP32 | | SLC6A11 | PNISR | BCL2L1 |
| LYZ2 | ARRB1 | | TNIK | RPL5 | CAMK2A |
| MARCKS | ATP1B2 | | | RPL7 | CASS4 |
| MDGA2 | ATP2B2 | | | RPL7A | CEP68 |
| NRP2 | BCAN | | | RPL13A | CSF1R |
| RPL23 | BCAR3 | | | RPL14 | CYFIP1 |
| RPS4X | BRSK2 | | | RPL21 | DIP2B |
| SEMA6A | CACNA1A | | | RPL23 | DNM2 |
| SPARC | CARM1 | | | RPL32 | EP300 |
| STAT3 | CDH2 | | | RPLP0 | EPN1 |
| TRIM9 | CDH4 | | | RPS3A1 | FES |
| UNC13C | CLMN | | | RPS4X | FRMD4B |
| | CNTN1 | | | RPS12 | FRY |
| | DAAM2 | | | RPS28 | FSCN1 |
| | DAG1 | | | RPSA | GRK2 |
| | DAGLA | | | STX16 | HCFC1 |
| | DCC | | | C1QB | HRH2 |
| | DCHS1 | | | C1QC | HS6ST1 |
| | DENND1A | | | CFL1 | ITPR2 |
| | EFNA5 | | | EEA1 | JADE2 |
| | EPHB1 | | | EIF5B | LIFR |
| | EPN1 | | | MYO9A | MGLL |
| | ETV4 | | | PABPC1 | MINK1 |
| | EYA1 | | | PNISR | NAV1 |
| | FAT1 | | | RPL5 | NOTCH2 |
| | FGFR1 | | | RPL7 | PRAG1 |
| | FMN2 | | | RPL7A | PREX1 |
| | GPR158 | | | RPL13A | PRKCD |
| | GFRA1 | | | RPL14 | PTPRO |
| | GLI2 | | | RPL21 | PTPRS |
| | GNAO1 | | | RPL23 | RAPGEF1 |
| | GRHL1 | | | RPL32 | RERE |
| | GRIN2C | | | RPLP0 | RGS10 |
| | HIPK2 | | | RPS3A1 | RIN3 |
| | KANK1 | | | RPS4X | SEMA4B |
| | KCNK2 | | | RPS12 | SIPA1L1 |
| | KIFC3 | | | RPS28 | SIPA1L3 |

| | | | | | |
|---|---|---|---|---|---|
| | MAPK8IP3 | | | RPSA | SMURF1 |
| | MINK1 | | | STX16 | SORT1 |
| | MPP7 | | | | TJP1 |
| | MTOR | | | | TNFRSF1B |
| | MTSS2 | | | | TRIO |
| | NEDD4L | | | | VAV1 |
| | NOTCH1 | | | | WASF2 |
| | NOTCH2 | | | | WHRN |
| | NR1D1 | | | | ZMIZ1 |
| | NRBP2 | | | | |
| | NTSR2 | | | | |
| | PCDHGC3 | | | | |
| | PDGFRB | | | | |
| | PLPP3 | | | | |
| | PLXNB1 | | | | |
| | PREX1 | | | | |
| | PSD2 | | | | |
| | PSAP | | | | |
| | PTPRS | | | | |
| | RGS6 | | | | |
| | SCRN1 | | | | |
| | SEPTIN5 | | | | |
| | SGIP1 | | | | |
| | SIPA1L3 | | | | |
| | SLC6A1 | | | | |
| | SLC7A10 | | | | |
| | SLC38A1 | | | | |
| | SORBS3 | | | | |
| | SORCS2 | | | | |
| | SOX1 | | | | |
| | SYNGAP1 | | | | |
| | TRAK1 | | | | |
| | VEGFA | | | | |
| | ZMIZ1 | | | | |

**Unique upregulated or downregulated genes associated with astrocytes or microglia from *SRR$^{fl/fl}$* and *Tmem119$^{creErt2}$:SRR$^{fl/fl}$* mice.** Consolidated gene list from all astrocytes or microglia subclusters show uniquely expressed upregulated or downregulated genes associated with *SRR$^{fl/fl}$* and *Tmem119$^{creErt2}$:SRR$^{fl/fl}$* mice.

**Table S5.**

| Sample ID | Protein or RNA | Reason | Region | GCS at Arrival | Time (from arrival to surgery) | Age | Sex | Race | CNS Co-morbidities |
|---|---|---|---|---|---|---|---|---|---|
| C1 | Protein | Cardiac arrest | CC | N/A | N/A | 22 | M | N/A | N/A |
| C2 | Protein | Cardiac arrest | CC | N/A | N/A | 66 | M | N/A | N/A |
| C3 | Protein | Cardiac arrest | F | N/A | N/A | 82 | M | N/A | N/A |
| C4 | Protein | Cardiac arrest | F | N/A | N/A | 22 | M | N/A | N/A |
| C5 | Protein | Cardiac arrest | T | N/A | N/A | 77 | M | N/A | N/A |
| C6 | Protein | Cardiac arrest | T | N/A | N/A | 66 | M | N/A | N/A |
| C7 | Protein | Cardiac arrest | P | N/A | N/A | 66 | M | N/A | N/A |
| C8 | Protein | Cardiac arrest | WB | N/A | N/A | 82 | M | N/A | N/A |
| C9 | RNA | Cardiac arrest | O | N/A | N/A | 22-68 | 14 M/F | W | N/A |
| C10 | RNA | Cardiac arrest | P | N/A | N/A | 35-89 | 4 M/F | W | N/A |
| C11 | RNA | Cardiac arrest | T | N/A | N/A | 21-82 | 5 M | A | N/A |
| TBI1 | Protein | Fall | F | 10 | <1 day | 32 | M | W | Alcohol use disorder |
| TBI2 | Both | GSW | P/O | 14 | 2 days | 21 | M | B | None |
| TBI3 | Protein | MVA | F | 7T | 6 days | 50 | M | W | None |
| TBI4 | Both | MVA | T | 13, rapidly deteriorating | 2 hours | 76 | M | B | Parkinson’s disease |
| TBI5 | Protein | Fall | F | 7T | 2 days | 54 | F | B | None |
| TBI6 | Both | GSW | T | 7T | 1-2 hours | 18 | M | B | None |
| TBI7 | Both | MVA | F | 10 | 5-6 hours | 63 | M | W | None |
| TBI8 | Protein | Fall | T | 6T | 15+ hours | 28 | F | W | Epilepsy |
| TBI9 | Both | Bike accident | F | 7 | 1 hour | 44 | M | H | None |
| TBI10 | Protein | MVA | T | 7T | 4 days | 34 | M | H | None |
| TBI11 | Both | MVA | T | 3 | 1 day | 58 | M | N/A | None |
| TBI12 | Both | MVA | F+T | 3 | 1 hour | 29 | M | H | None |
| TBI13 | Both | Bike accident | T | 11 | 1 day | 67 | M | W | None |

| | | | | | | | | | |
|---|---|---|---|---|---|---|---|---|---|
| TBI14 | Both | MVA | F | 3 | 1 hour | 22 | M | B | None |
| TBI15 | Both | Fall | F | 5 | NA | 78 | M | H | Frontotemporal dementia, aphasia |
| TBI16 | RNA | MVA | CB | 13 | <1 day | 31 | M | W | Bipolar disorder, manic depression, substance use |
| TBI17 | RNA | MVA | F | 3 | 12 hours | 50 | F | W | Depression, anxiety, PTSD |
| TBI18 | RNA | Fall | F | 7 | 1-2 hours | 55 | M | H | Epilepsy |
| TBI20 | RNA | Fall | F | 5 | 4 hours | 27 | F | B | N/A |
| TBI21 | RNA | MVA | T | 3 | 6 hours | 27 | M | H | None |
| TBI22 | RNA | NA | NA | NA | NA | 60 | M | NA | Substance use disorder |
| TBI23 | RNA | NA | T | 4 | 6 hours | 70 | M | H | Alcohol use disorder |
| TBI24 | RNA | Fall | F | 15 | 4 days | 43 | M | H | None |
| TBI25 | RNA | Assault | F | 3 | 11 hours | 47 | M | W | Substance use disorder |
| TBI26 | RNA | GSW | F | 15 | 7 hours | 62 | M | B | None |
| TBI27 | RNA | MVA | F | 3 | 5 days | 37 | M | H | None |
| TBI28 | RNA | GSW | NA | 10 | 3 hours | 37 | M | H | None |
| TBI29 | RNA | MVA | NA | 4 | 3 hours | 29 | M | H | ADHD, depression, schizophrenia |
| TBI30 | RNA | Fall | T | 7 | 5 hours | 85 | F | W | Depression |
| TBI31 | RNA | Assault | P | 14 | 6 hours | 55 | M | W | None |
| TBI32 | RNA | Assault | P | 4 | <1 hour | 60 | M | H | None |
| TBI33 | RNA | Fall | T | 15 | 8 hours | 74 | F | H | Epilepsy, mood disorder |
| TBI34 | RNA | MVA | NA | 3 | 22 hours | 28 | M | H | None |
| TBI35 | RNA | GSW | T | 7 | 9 hours | 23 | M | W | None |
| TBI36 | RNA | GSW | NA | 10 | NA | 20 | M | B | None |
| TBI37 | RNA | MVA | NA | 3 | NA | 28 | M | H | None |

**Injury and demographic information on human brain samples.** A total of 12 TBI patients were processed for Western blot and qPCR analysis. C1-C11 reference control samples. MVA = motor vehicle accident, GSW = gunshot wound, WB = whole brain, CC = cerebral cortex, O = occipital, P = parietal, F = frontal, T = temporal, CB = cerebellum, M = male, F = female, B = Black, W = white, A = Asian, H = Hispanic. GCS = Glasgow Coma Scale, where a score of 1-8 is classified as a severe injury, 9-12 as moderate, and 13-15 as mild; T indicates the patient is intubated.

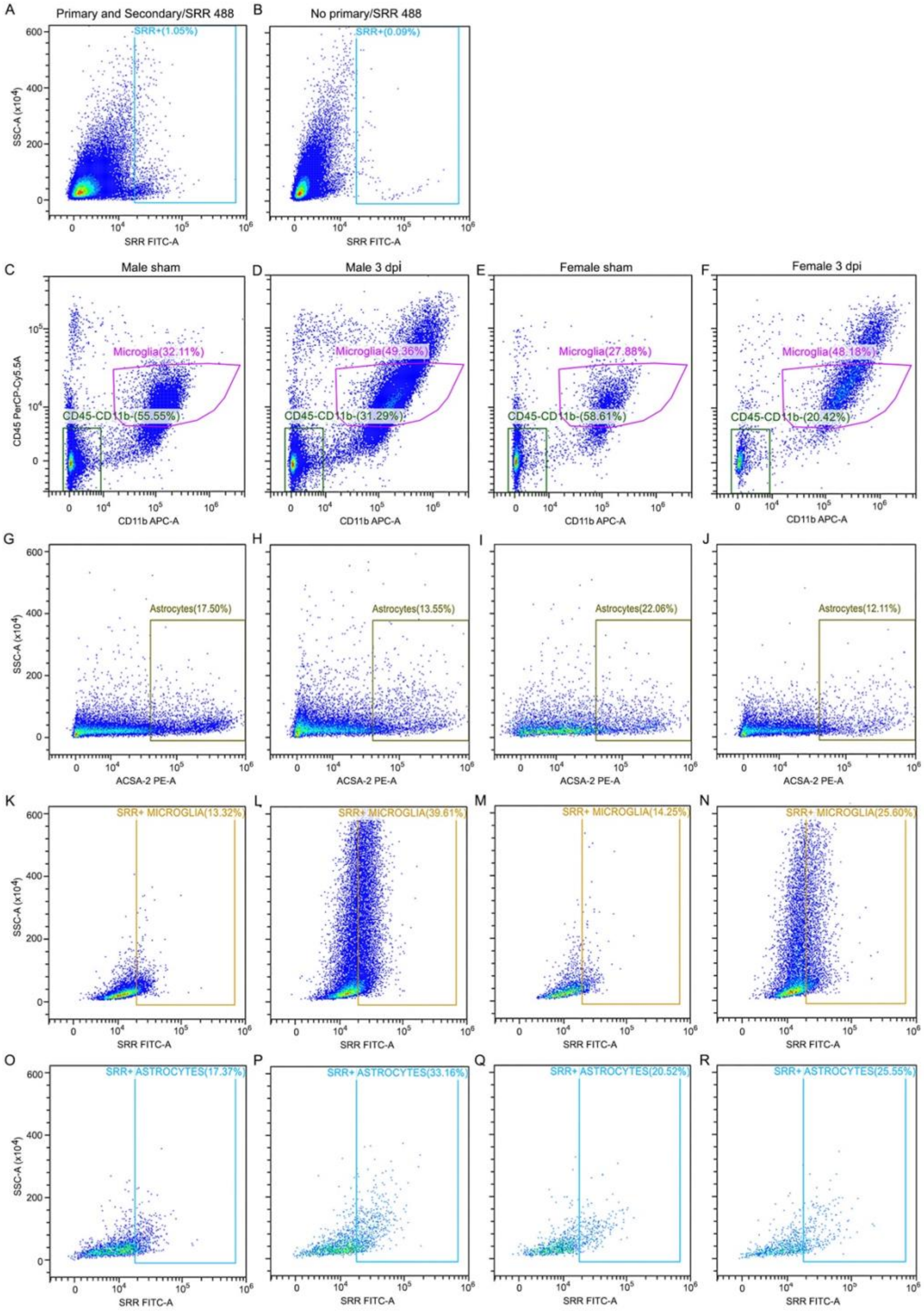

**Figure S1. Representative plots illustrating gating strategy for SRR flow cytometry.** (A) SRR antibody gating was determined based on (B) secondary antibody only staining. (C-F) $CD45^{Low}$ and CD11b+ microglia cell populations were defined from DAPI-labeled viable cells, and (G-J) ACSA-2+ astrocytes were defined from remaining CD45-, CD11b- cells from male and female sham and CCI-injured hippocampi. Gating strategy for SRR+ (K-N) microglia and (O-R) astrocytes.

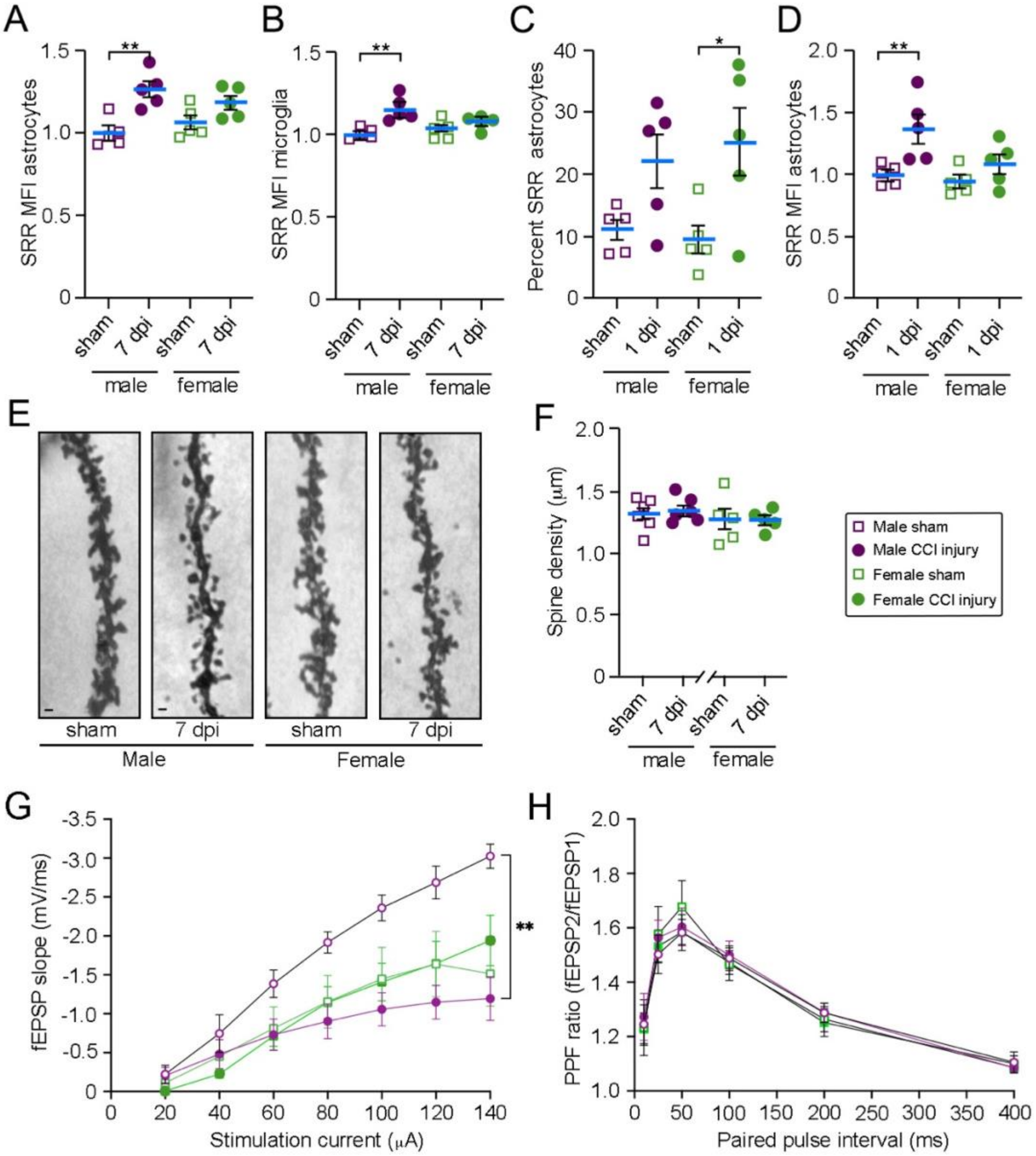

**Figure S2. Female glia cells express less SRR per cell than male glia after CCI injury.** (A-B) Flow cytometry analysis shows significantly increased mean fluorescent intensity (MFI) per SRR positive hippocampal astrocyte and microglia cell in male but not female mice at 7 dpi. (C) Female hippocampal astrocytes significantly upregulate SRR at 1 dpi; however, similar trends are observed in males. (D) The MFI per astrocyte is significantly increased only in male mice. (E) Representative images of Golgi staining in the dendrites of pyramidal cells in the CA1 hippocampus, where (F) dendritic spine density is unchanged in WT mice after injury. (G) Input/output electrophysiology curves. Scale bar: 1 μM. *p<0.05, **p<0.01. (A-D) Two-way ANOVA with Tukey's multiple comparison test; (G,H) Mixed -effects model with Tukey multiple comparisons test. (A-D) n=5, (F) n=5-6, (G-H) n=5-7. Values represent mean ± SEM.

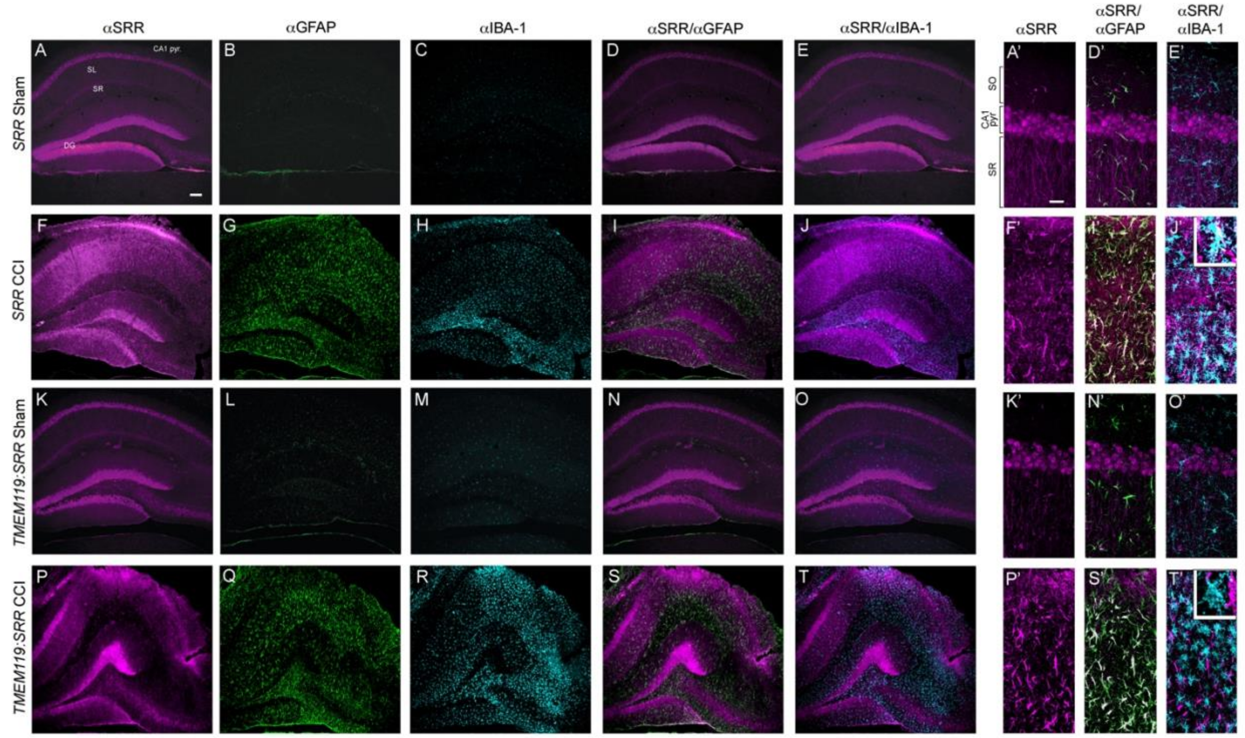

**Figure S3. SRR is removed from microglia cells following tamoxifen treatment in female mice.** (A-E, A’,D’,E’) SRR expression follows a predominantly neuronal expression pattern in sham conditions in $SRR^{fl/fl}$ control and (K-O, K’-O’) $Tmem119^{creERT2}$:$SRR^{fl/fl}$ female mice at 7 dpi. (F-J, F’,I’,J’, P-T, P’-T’) CCI-injury induces increased gliosis and glial SRR expression, where SRR is co-localized with (S’) astrocytes but not (T’) microglia in $Tmem119^{creERT2}$:$SRR^{fl/fl}$ mice.

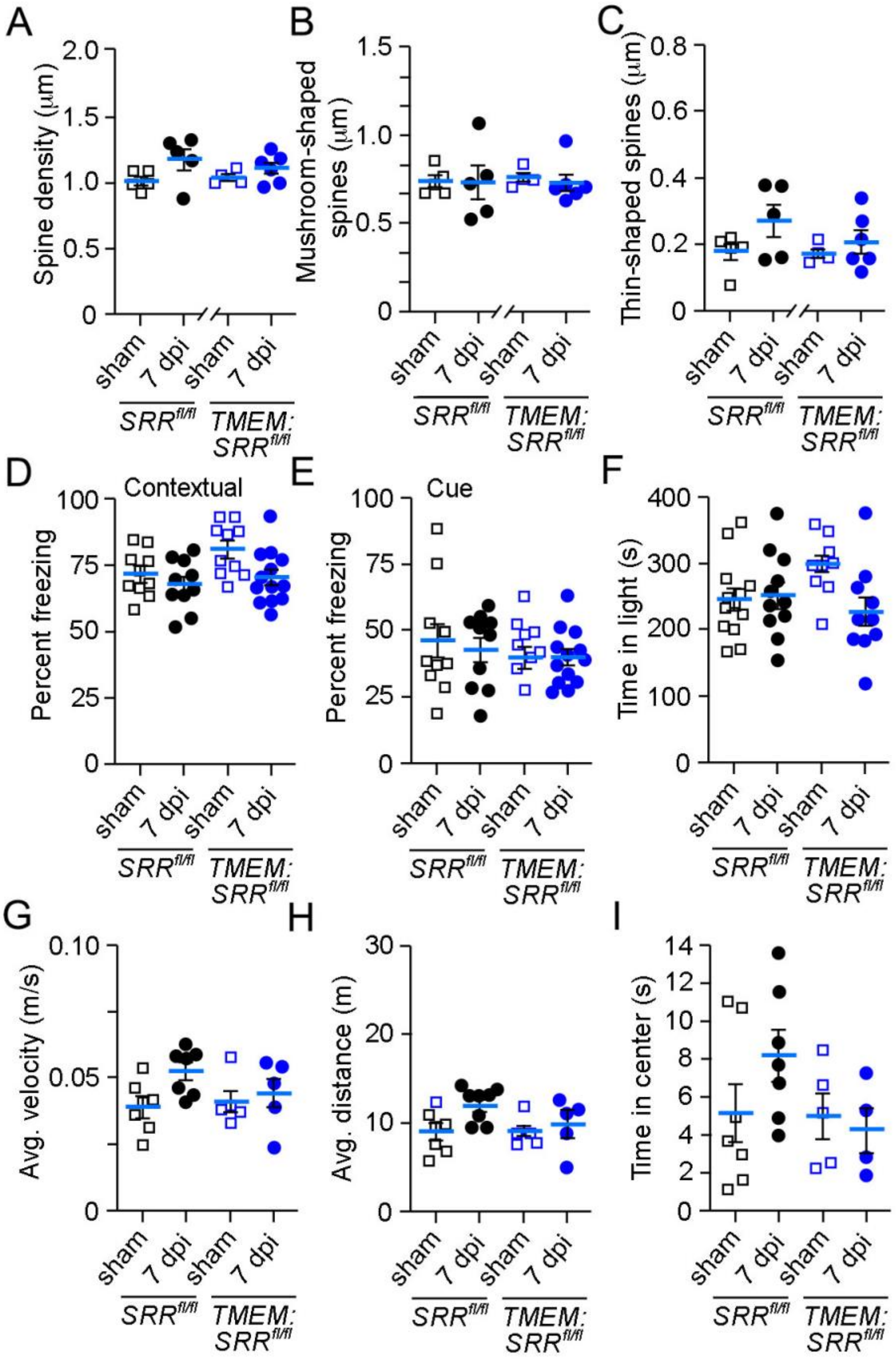

**Figure S4. CCI injury or microglial D-serine does not affect dendritic spine damage, contextual fear conditioning, anxiety, or locomotion in female mice.** (A-C) Golgi analysis shows no change in dendritic spine density or morphology in female *SRR*$^{fl/fl}$ or *Tmem119*$^{creERT2}$:*SRR*$^{fl/fl}$ mice at 7 dpi. (D) No deficit in contextual or (E) cued fear conditioning behavior in female mice at 7 dpi. (F) No significant difference in anxiety-like behavior in the light-dark transition test or (I) open-field assay, as well as no changes in the (G) average velocity or (H) distance travelled during open-field in female mice at 7 dpi. Values represent mean ± SEM.

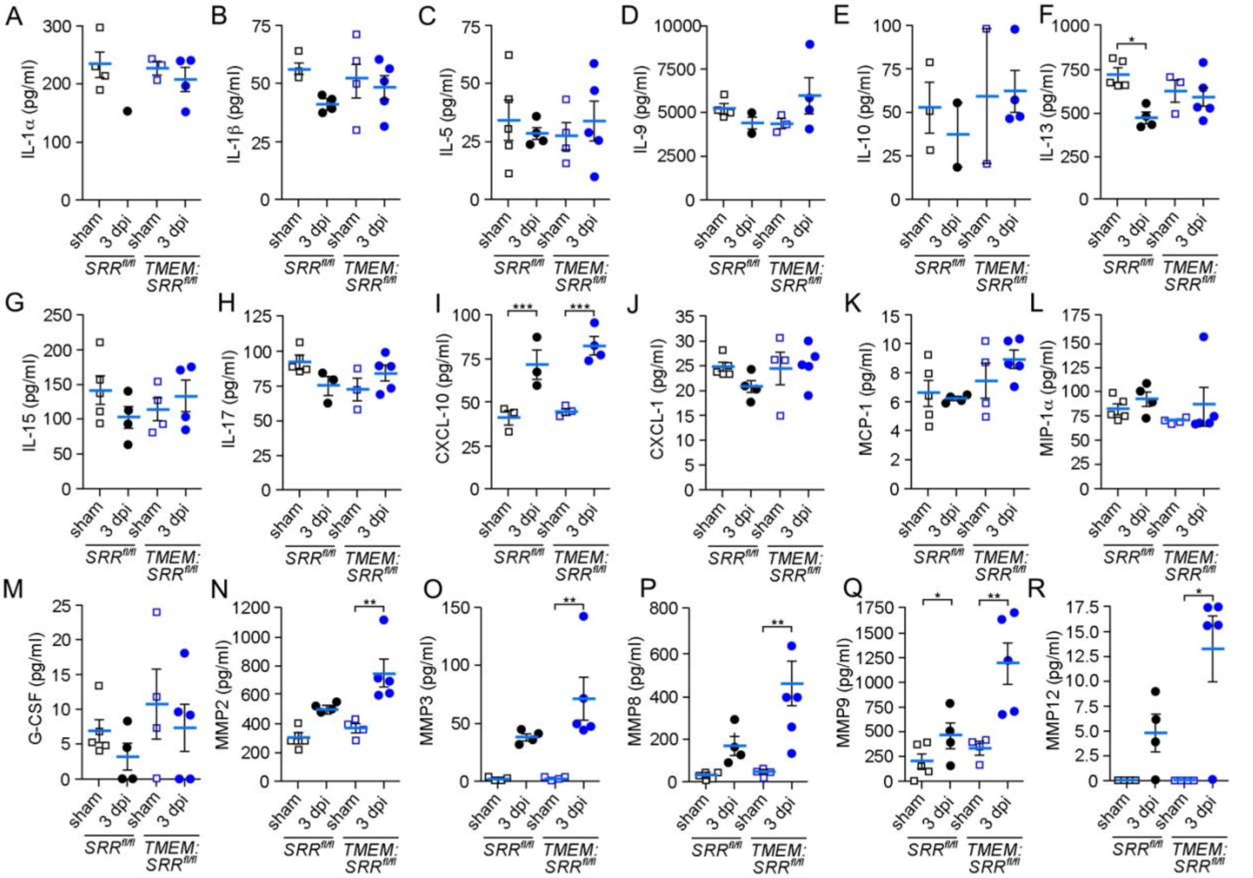

**Figure S5. Microglial SRR ablation affects matrix metalloproteinases (MMPs) and IL-13 expression after injury.** (A-H) No significant changes interleukin (IL)-1a, IL-1b, IL-5, IL-9, IL-10, IL-15, or IL-17 in either genotype at 3 dpi. (F) IL-13 is significantly downregulated at 3 dpi in control but not microglial SRR knockout mice. (I) The chemokine CXCL-10 is significantly increased after injury, (J-M) whereas chemokines CXCL-1, MCP-1, MIP-1a, and G-CSF are unaffected. (N-R) CCI-injury induces significant upregulation of MMP2, MMP3, MMP8, MMP9, and MMP12 in *Tmem119*$^{cre:ERT2}$*:SRR*$^{fl/fl}$ male mice at 3 dpi. (Q) MMP9 expression is significantly higher after injury in *Tmem119*$^{cre:ERT2}$*:SRR*$^{fl/fl}$ than *SRR*$^{fl/fl}$ mice. *p<0.05, **p<0.01, ***p<0.001. Two-way ANOVA with Tukey's multiple comparison test; n=1-5. Values represent mean ± SEM.

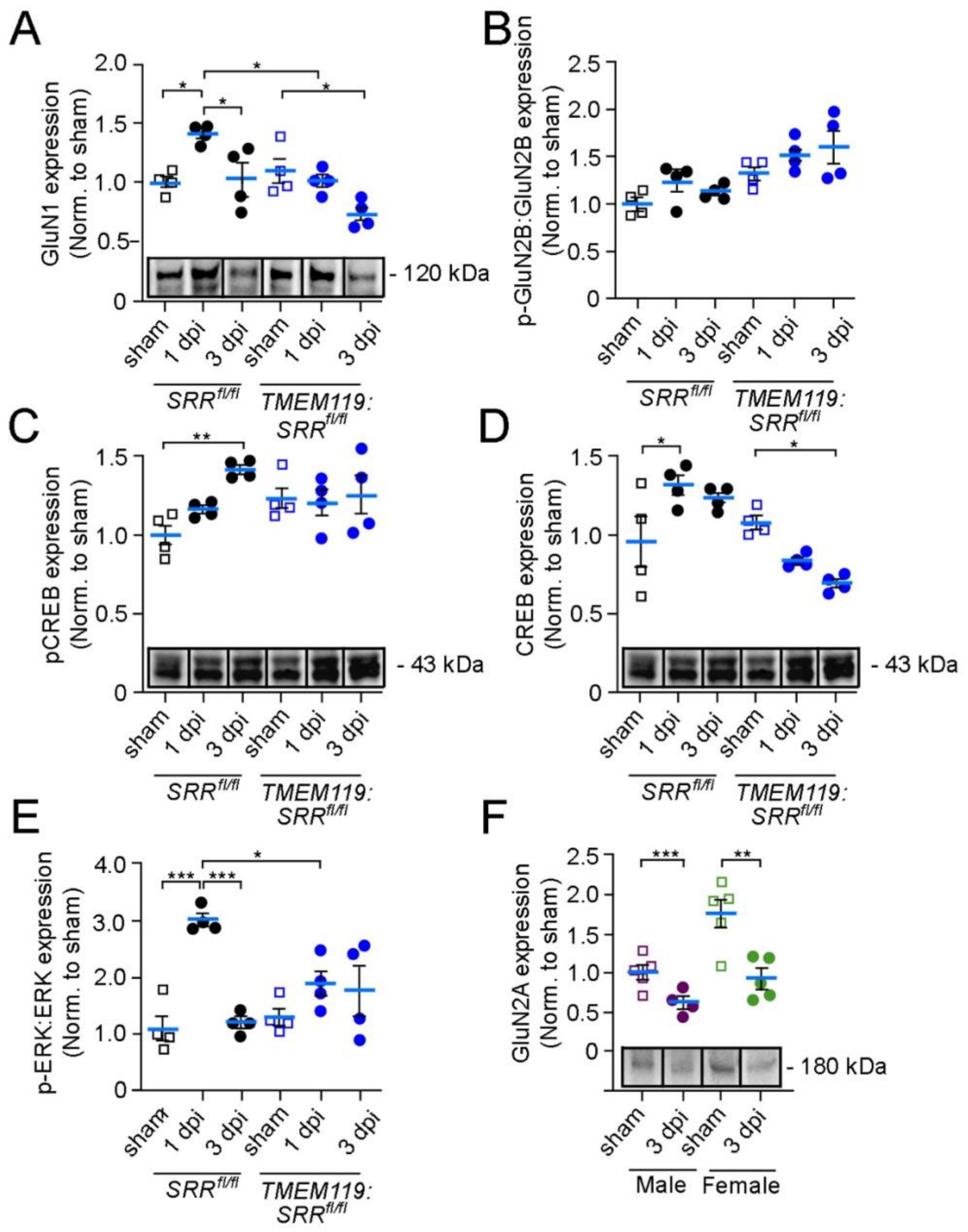

**Figure S6. Alterations in hippocampal synaptic proteins after CCI injury.** (A) Western blot of whole hippocampus shows a significant increase in GluN1 expression at 1 and 3 dpi in *SRR*$^{fl/fl}$ mice and decrease in *TMEM119*$^{creERT2}$*:SRR*$^{fl/fl}$ mice by 3 dpi. (B) There are no significant changes in the ratio of phosphorylated GluN2B to GluN2B after injury. (C) There is a significant increase in phosphorylated CREB expression normalized to total protein in *SRR*$^{fl/fl}$ mice at 3 dpi, as well as (D) an increase in total CREB expression at 1 dpi in *SRR*$^{fl/fl}$ mice and decrease by 3 dpi in *Tmem119*$^{creERT2}$*:SRR*$^{fl/fl}$ mice. (E) The ratio of phosphorylated to non-phosphorylated ERK1/2 is significantly increased in control but not microglial *SRR*$^{fl/fl}$ mice at 1 dpi. (F) GluN2A expression is significantly downregulated in isolated hippocampal membrane fraction in WT female and male mice at 3 dpi. *$p<0.05$, **$p<0.01$, ***$p<0.001$. Two-way ANOVA with Tukey's multiple comparison test; n=4-5. Values represent mean ± SEM.

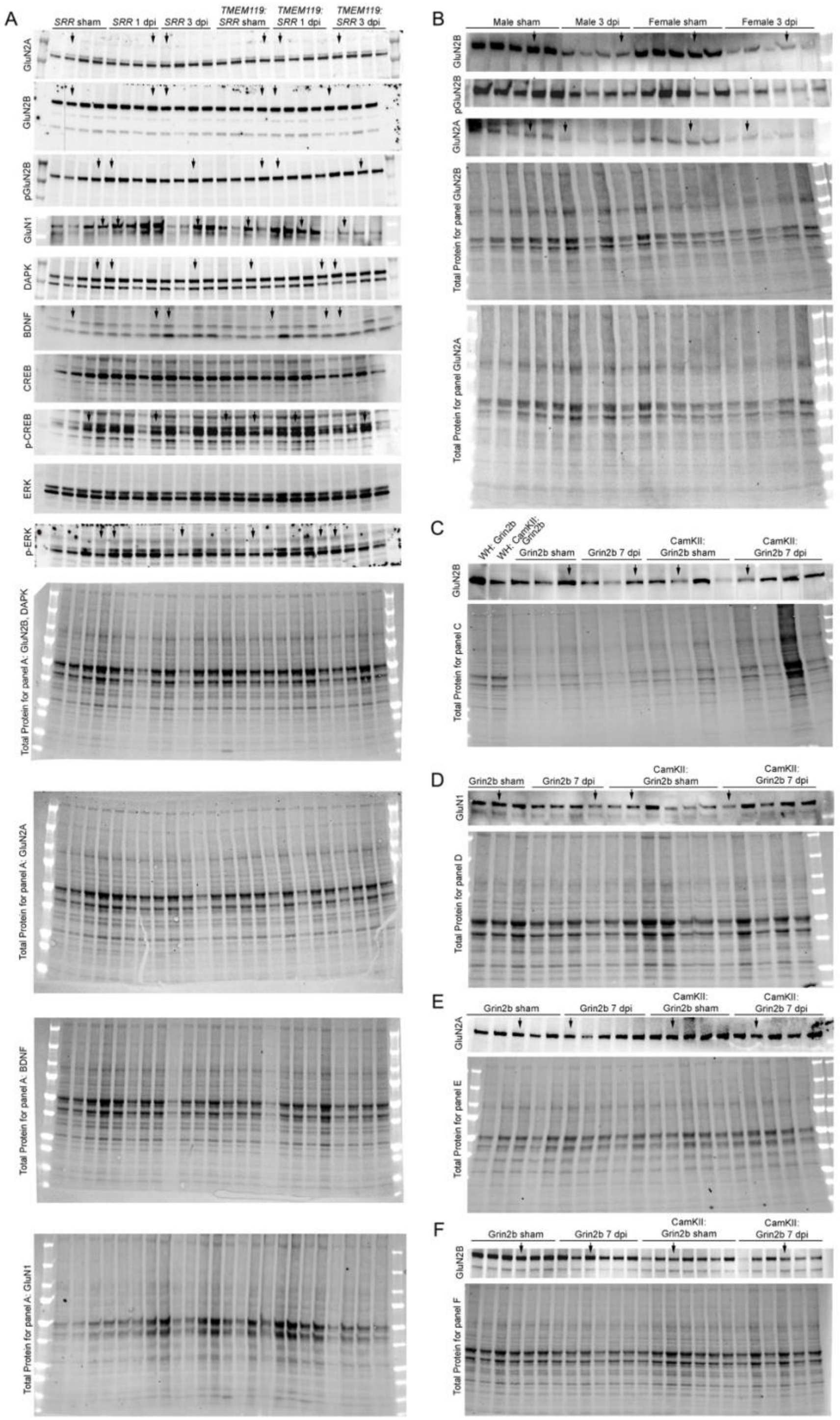

**Figure S7. Uncut and unaltered Western blot images.** Images were used for densitometry analysis in Figures (A,B) 2, S6, (C) 3, and (D-F) S8, where proteins of interest were normalized to total protein. (A,D-F) Samples were isolated from whole hippocampal lysate and (B,C) isolated plasma membrane fraction. WH = whole hippocampus.

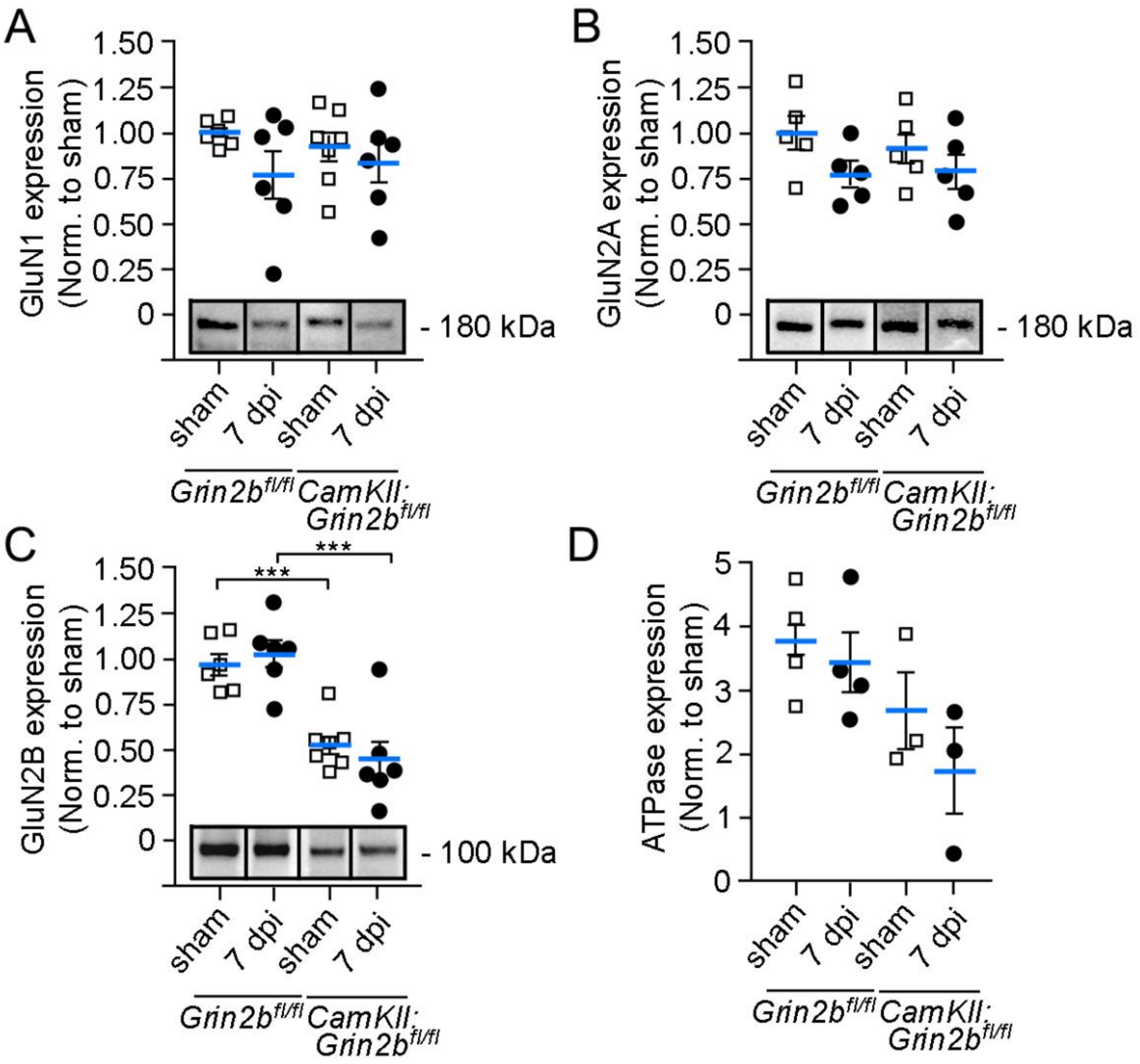

**Figure S8. Specific ablation of the GluN2B subunit of NMDARs in *CamkII$^{creERT2}$:Grin2b$^{fl/fl}$* mice.** (A) Western blot analysis of whole hippocampal tissue shows that expression levels of GluN1 and (B) GluN2A NMDAR subunits are unchanged in *CamKII$^{creERT2}$:Grin2b$^{fl/fl}$* as compared to control *Grin2b$^{fl/fl}$* sham or CCI male mice. (C) There is a significant reduction in the GluN2B subunit in whole hippocampal tissue in sham and CCI *CamKII$^{creERT2}$:Grin2b$^{fl/fl}$* mice. (D) Expression of the plasma membrane Na+/K+ ATPase is enriched in lysates from isolated hippocampal membrane fraction relative to whole hippocampal tissue. ***$p<0.001$. Two-way ANOVA with Tukey's multiple comparison test; n=4-7. Values represent mean ± SEM.

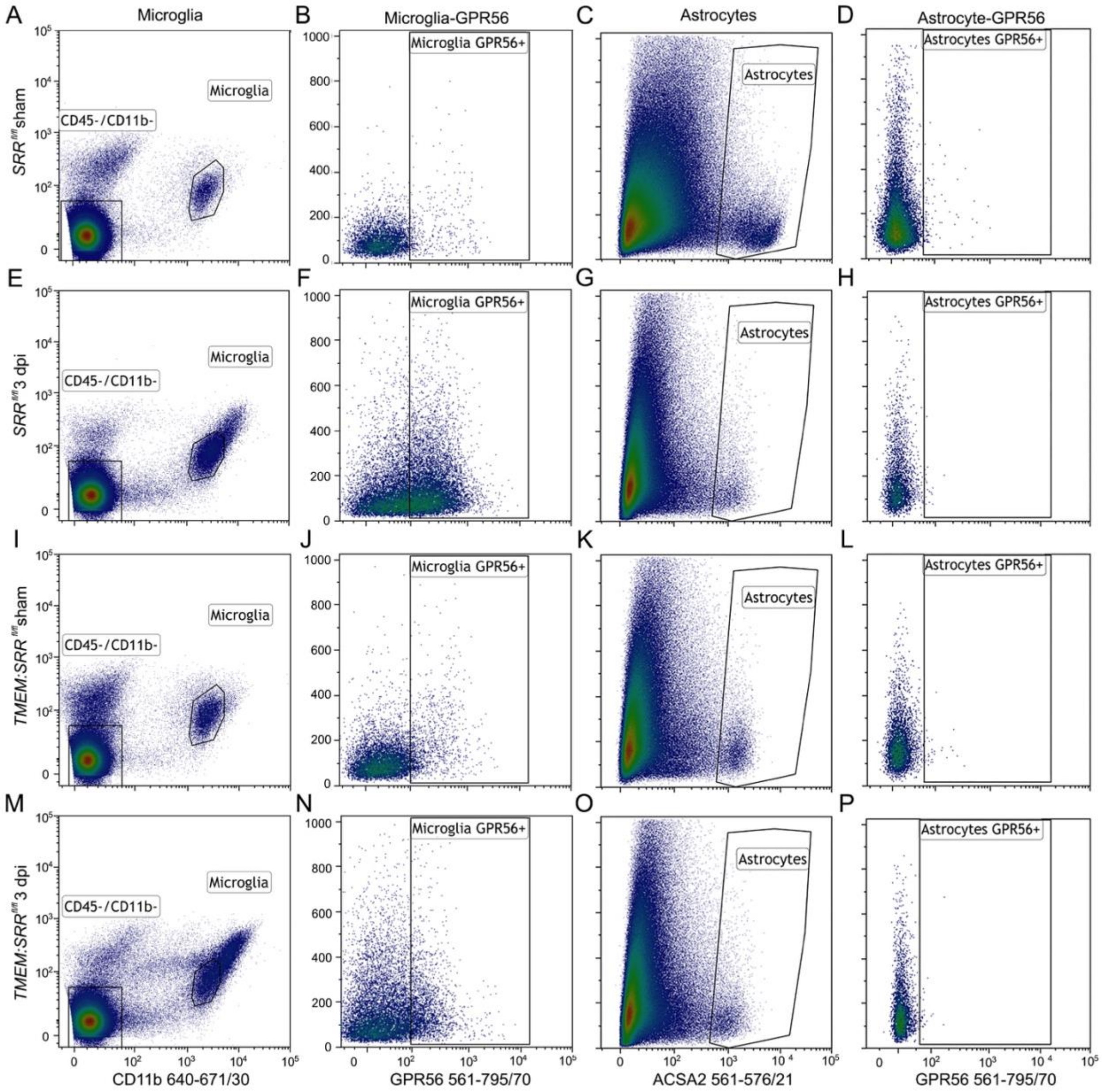

**Figure S9. Representative plots illustrating gating strategy for GPR56 in FACS microglia and astrocytes.** (A,E,I,M) CD45$^{Low}$ and CD11b+ microglia cell populations were defined from DAPI-labeled viable cells, and (C,G,K,O) ACSA-2+ astrocytes were defined from CD45-, CD11b- DAPI-labeled viable hippocampal cells. (B,J) Low levels of GPR56 in sham microglia are increased in 3 dpi microglia isolated from (F) *SRR*$^{fl/fl}$ and to a lesser extent in (N) *Tmem119*$^{creERT2}$*:SRR*$^{fl/fl}$ mice. (D,H,L,P) Low GPR56 expression was detected in hippocampal sham or 3 dpi astrocytes.

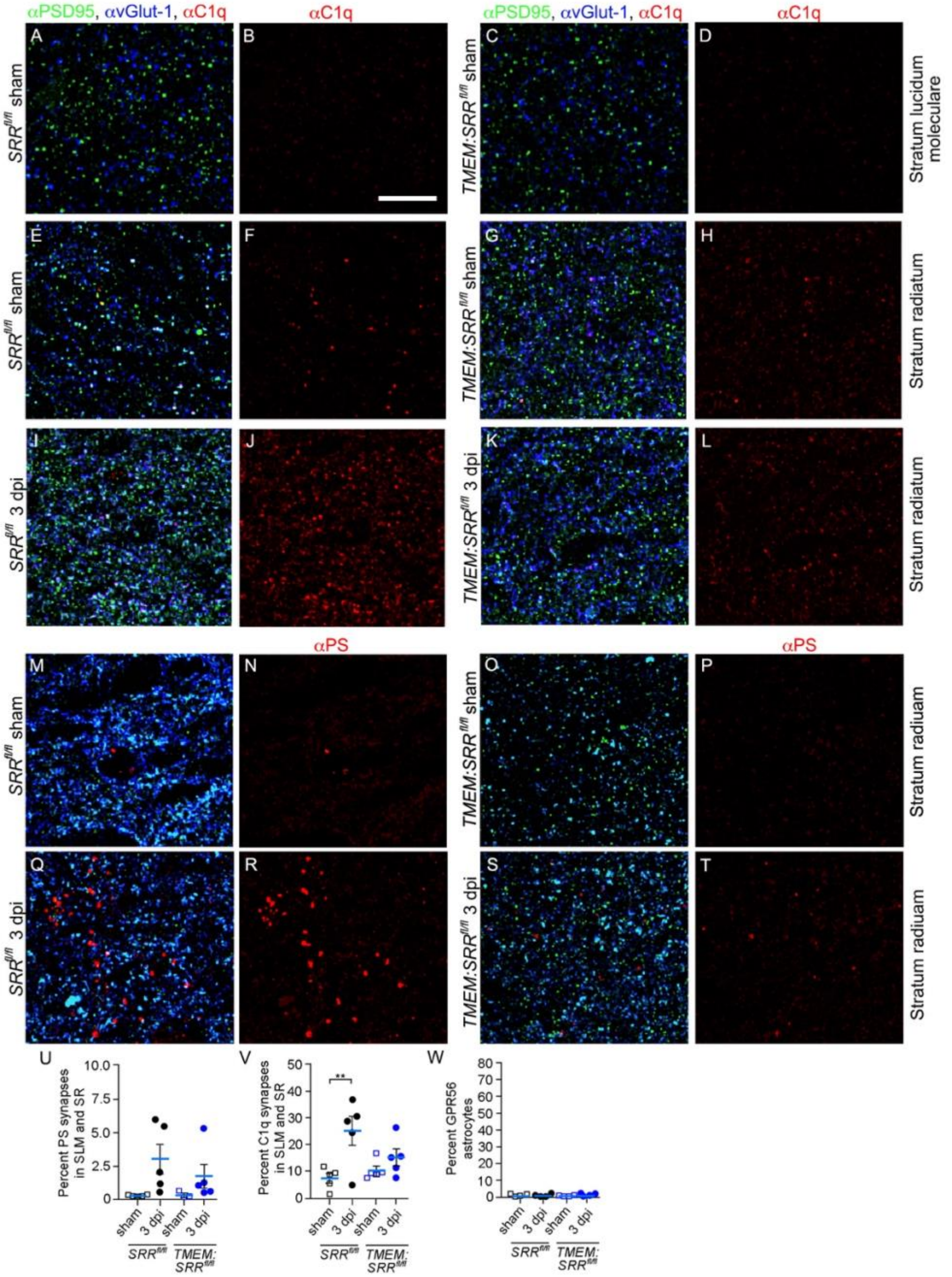

**Figure S10. PS and C1q association with synapses in the *stratum lucidum moleculare (SLM)* and *stratum radiatum (SR)* after injury.** (A-D) C1q is expression is low in the *SLM* and (E-H) *SR* of the CA1 hippocampus in sham conditions but is (I-L) upregulated at 3 dpi in both $SRR^{fl/fl}$ and $Tmem119^{creERT2}$:$SRR^{fl/fl}$ mice in the *SR*. (M-P) Similarly, PS expression is low in the (M-P) *SR* under sham conditions and (Q-T) upregulated at 3 dpi. (U) Combined quantification of PS+ synapses in the *SLM* and *SR* shows no significant differences after injury, whereas (V) C1q+ synapses are significantly upregulated at 3 dpi in $SRR^{fl/fl}$ mice. (W) Hippocampal FACS astrocytes show low levels of GPR56 expression that are unaffected by CCI-injury. Scale bar: 5 μM. **p<0.01. Two-way ANOVA with Tukey's multiple comparison test; n=3-5 mice/group each with 3 sections/area. Values represent mean ± SEM.

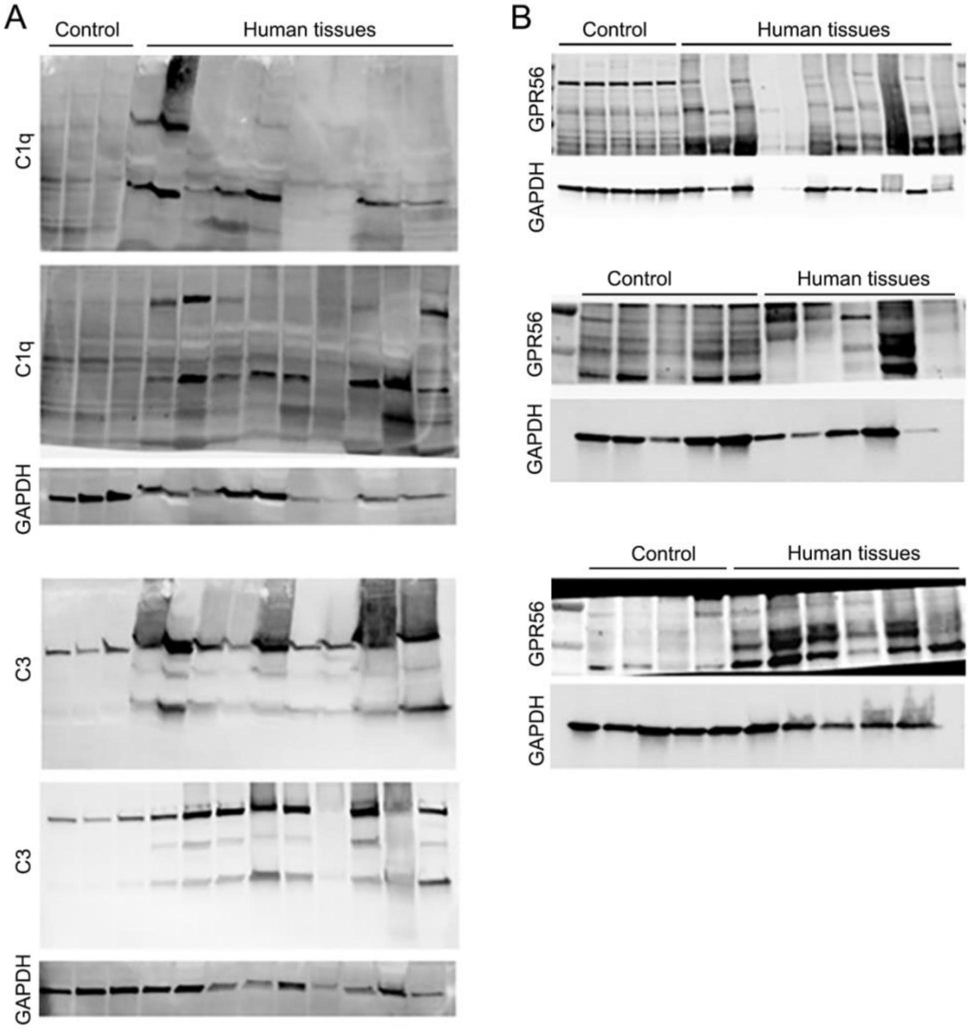

**Figure S11. Uncut and unaltered Western blot images.** Images were used for densitometry in Figure 7, where proteins of interest were normalized to GAPDH.

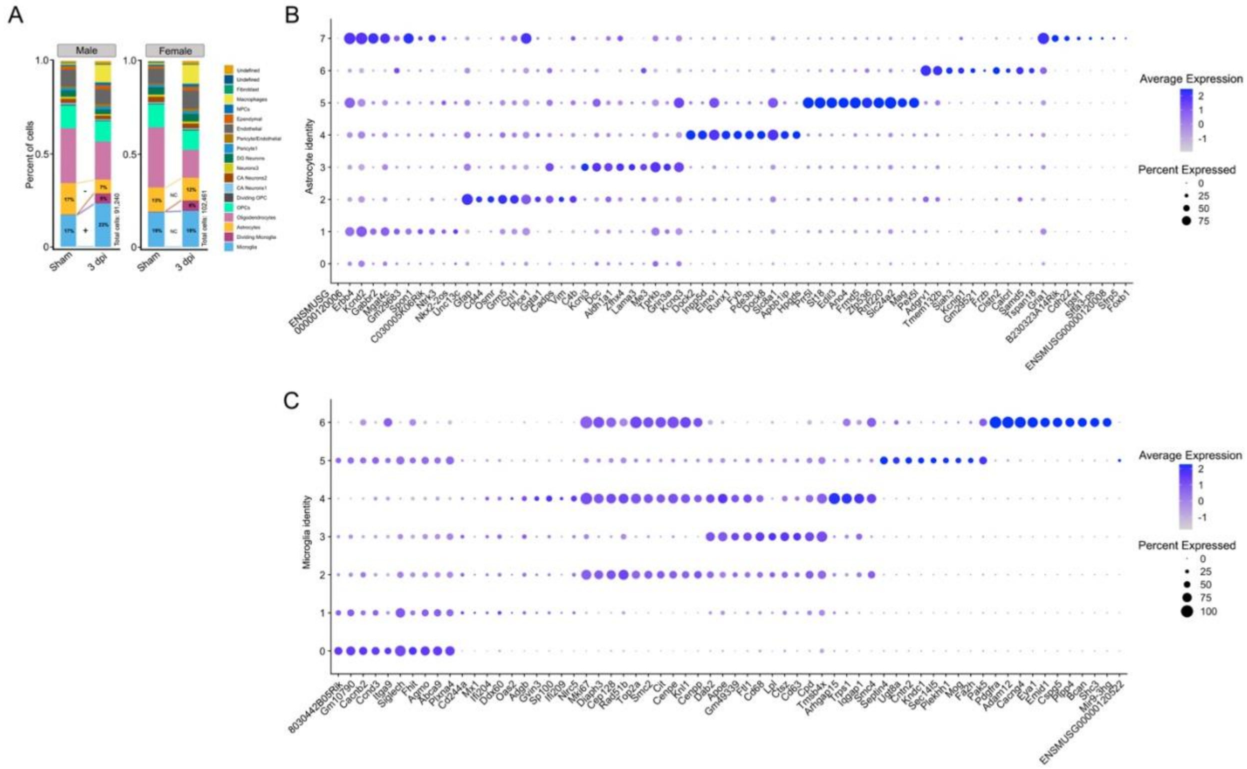

**Figure S12. Cell distribution and categorization of astrocyte and microglia subclusters in male and female mice.** (A) Percent of cell populations were determined for male and female mice that show increased (+) or decreased (-) percentages between sham and CCI injured groups. (B) Top 10 marker genes for 8 astrocyte subclusters and 7 microglia subclusters.

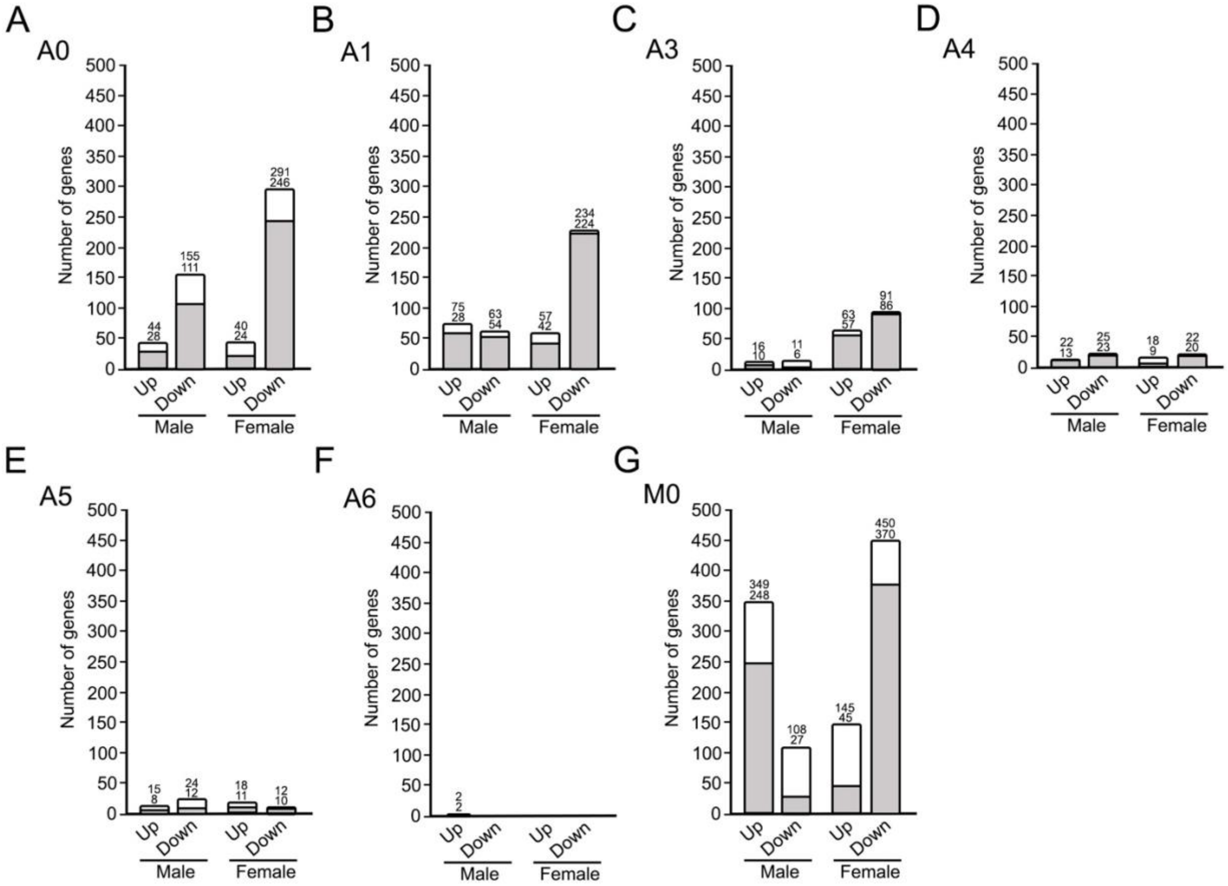

**Figure S13. Comparison of astrocyte and microglia subclusters that were present in both sham and CCI injured male and female mice.** (A-G) Total (open bars) and unique genes (gray bars) are shown for all subclusters that are present in sham and CCI injury, where significantly upregulated and downregulated genes are shown. The term unique is defined as genes unique to males or females between the up- or down-regulated groups.

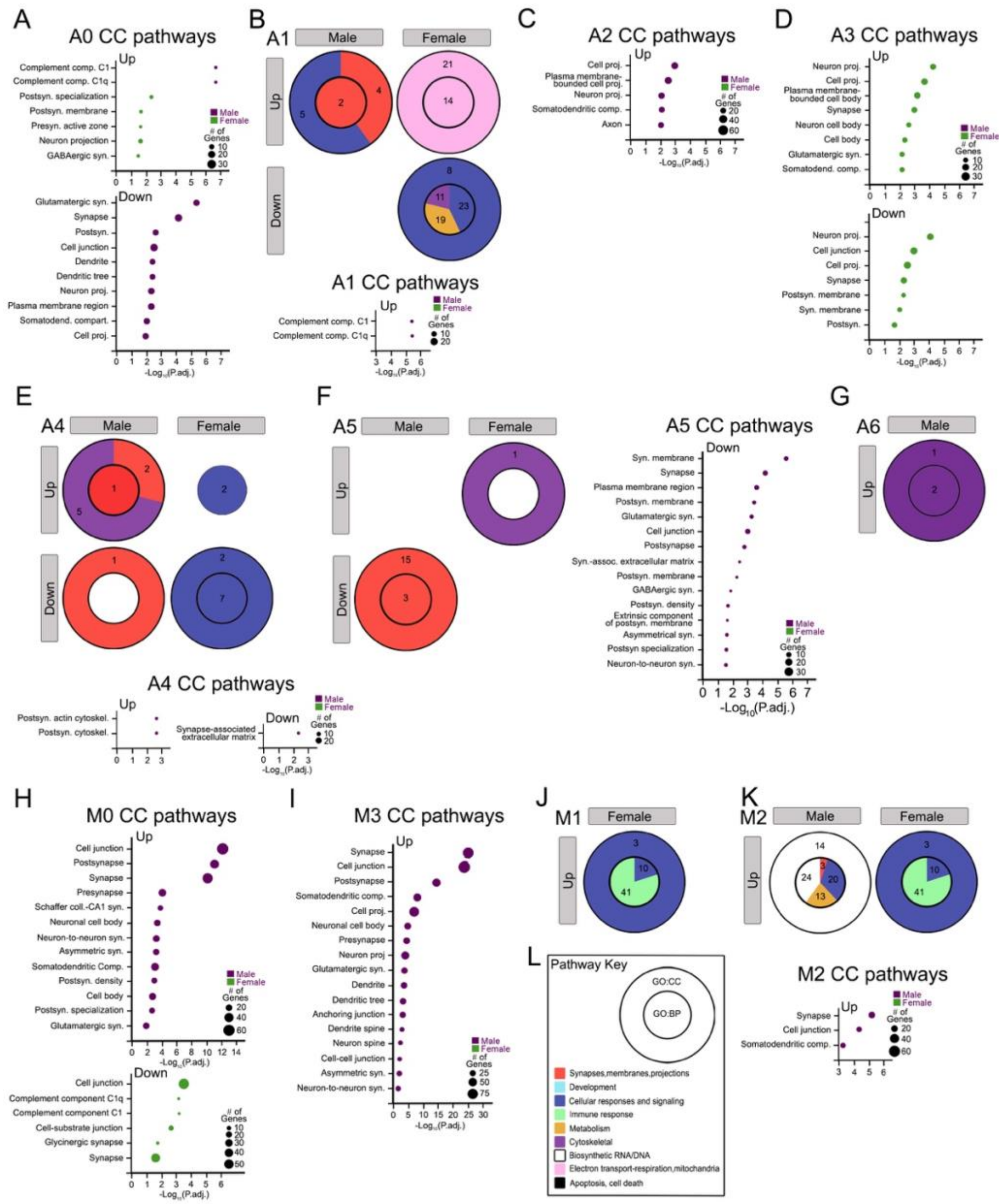

**Figure S14. Pathway analysis illustrates changes in synaptic-related pathways for astrocyte and microglia subclusters in male and female mice**. (A) Astrocyte subcluster A0, (B) A1, (D) A3, (E) A4, and (F) A5 pathway analysis using dot (i.e. bubble) plots for upregulated and downregulated GO:CC pathways in CCI relative to sham conditions. (C) Astrocyte subcluster A2 pathway analysis using bubble plots for upregulated GO:CC pathways, where A2 subclusters were comparisons between CCI male and CCI female mice. (G) Astrocyte subcluster A6 pathway analysis using pie plots. (H) Microglia subcluster M0 pathway analysis using bubble plots for upregulated and downregulated GO:CC pathways in CCI relative to sham conditions. (I) Microglia subcluster M3, (J) M1, and (K) M2 pathway analysis using bubble plots for upregulated GO:CC pathways between CCI male and CCI female mice. (L) Pathway key showing the cellular component (CC) in the outer pie and biological processes (BP) in the inner pie, where consolidated pathways are color-coded. Numbers in each pie region represents the number of consolidated GO pathways.

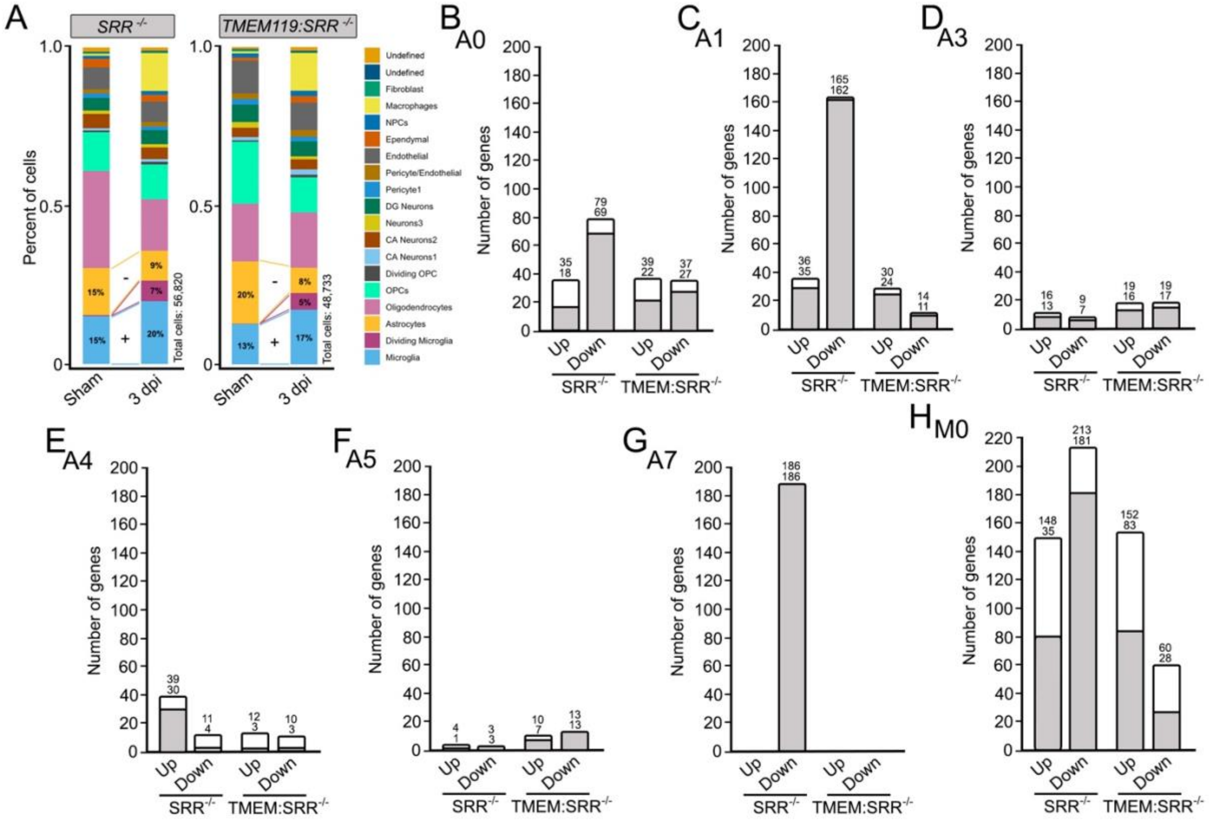

**Figure S15. Cell distribution and comparison of astrocyte and microglia subclusters that were present in both sham and CCI injured *SRR*$^{fl/fl}$ and *Tmem119*$^{creERT2}$*:SRR*$^{fl/fl}$ mice.** (A) Percent of cell populations were determined for *SRR*$^{fl/fl}$ and *Tmem119*$^{creERT2}$*:SRR*$^{fl/fl}$ mice that show increased (+) or decreased (-) percentages between sham and CCI injured groups. (B-H) Total (open bars) and unique genes (gray bars) are shown for all subclusters that are present in sham and CCI injury, where significantly upregulated and downregulated genes are shown. The term unique is defined as genes unique to *SRR*$^{fl/fl}$ and *Tmem119*$^{creERT2}$*:SRR*$^{fl/fl}$ mice between the up- or down-regulated groups.

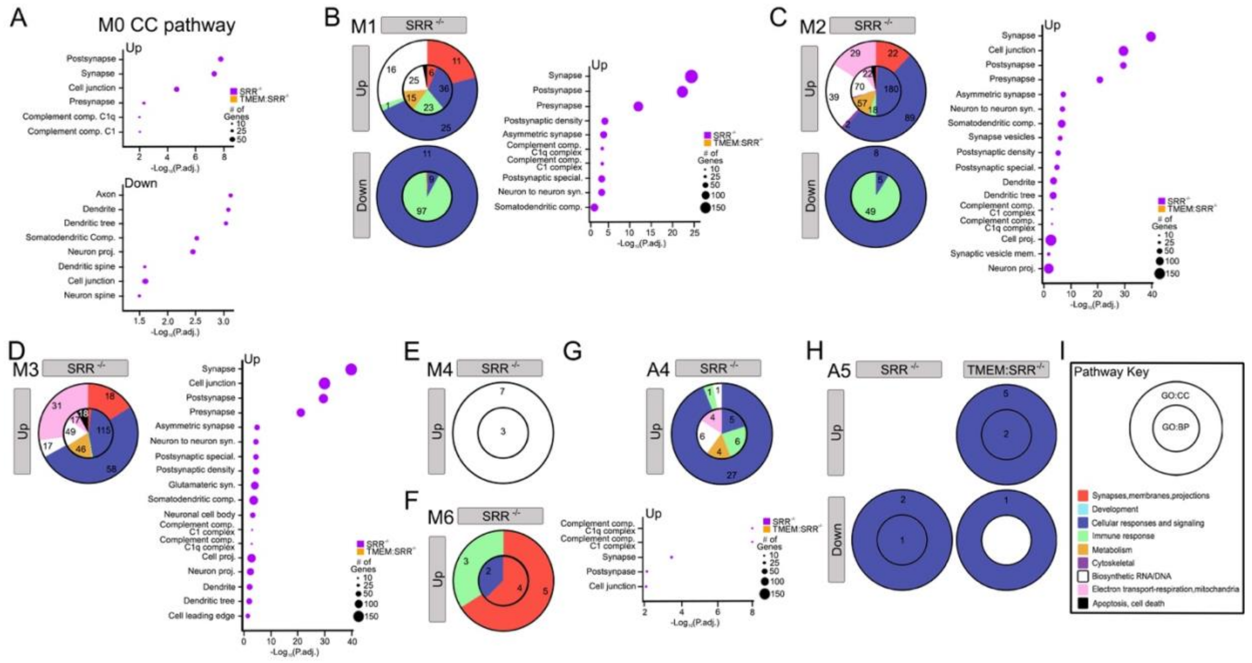

**Figure S16. Pathway analysis illustrates changes in synaptic-related pathways for astrocyte and microglia subclusters in *$SRR^{fl/fl}$* and *$Tmem119^{creERT2}$:$SRR^{fl/fl}$* mice**. (A) Microglia subcluster M0 pathway analysis using dot (i.e. bubble) plots for upregulated and downregulated GO:CC pathways in CCI relative to sham conditions. (B) Microglia subcluster M1, (C) M2, (D) M3, (E) M4, and (F) M6 pathway analysis using pie and bubble plots for upregulated GO pathways, where M1 subclusters were comparisons between CCI *$SRR^{fl/f}$* and CCI *$Tmem119^{creERT2}$:$SRR^{fl/f}$* mice. (G) Astrocyte subcluster A4 pathway analysis using pie and bubble plots for upregulated GO:CC pathways in CCI relative to sham conditions. (I) Pathway key showing the cellular component (CC) in the outer pie and biological processes (BP) in the inner pie, where consolidated pathways are color-coded. Numbers in each pie region represents the number of consolidated GO pathways.
